# High-fidelity, Large-scale Targeted Profiling of Microsatellites

**DOI:** 10.1101/2023.11.28.569106

**Authors:** Caitlin A. Loh, Danielle A. Shields, Adam Schwing, Gilad D. Evrony

## Abstract

Microsatellites are highly mutable sequences that can serve as markers for relationships among individuals or cells within a population. The accuracy and resolution of reconstructing these relationships depends on the fidelity of microsatellite profiling and the number of microsatellites profiled. However, current methods for targeted profiling of microsatellites incur significant “stutter” artifacts that interfere with accurate genotyping, and sequencing costs preclude whole-genome microsatellite profiling of a large number of samples. We developed a novel method for accurate and cost-effective targeted profiling of a panel of > 150,000 microsatellites per sample, along with a computational tool for designing large-scale microsatellite panels. Our method addresses the greatest challenge for microsatellite profiling — “stutter” artifacts — with a low-temperature hybridization capture that significantly reduces these artifacts. We also developed a computational tool for accurate genotyping of the resulting microsatellite sequencing data that uses an ensemble approach integrating three microsatellite genotyping tools, which we optimize by analysis of de novo microsatellite mutations in human trios. Altogether, our suite of experimental and computational tools enables high-fidelity, large-scale profiling of microsatellites, which may find utility in diverse applications such as lineage tracing, population genetics, ecology, and forensics.

## BACKGROUND

Microsatellites, also known as short tandem repeats, are genomic sequences comprised of tandem repeats of short (1 to 6 base pairs, bp) sequence motifs. Microsatellite loci vary widely in length and number of repeats, and there are more than one million microsatellite loci in the human genome that together comprise approximately 3% of the genome. The repetitive structure of microsatellites makes them highly mutable in comparison to other types of genomic sequences. The estimated de novo mutation rate of microsatellites is 1 × 10^−4^ to 1 × 10^−3^ per locus per generation, while the estimated de novo base substitution rate is 1.2 × 10^−8^ per base pair per generation (Sun et al. 2012; Kessler et al. 2020). Based on these de novo mutation rates, it is estimated that every cell division incurs several microsatellite mutations and that “silent” cell divisions with no microsatellite mutations are infrequent (Frumkin et al. 2005). This elevated mutability makes microsatellites highly variable within populations, and consequently, they are used extensively as markers of individuals members of a population in the fields of ecology (Selkoe and Toonen 2006; De Barba et al. 2017), forensics (Butler 2006; Moretti et al. 2016), and population genetics (Bruford and Wayne 1993; Putman and Carbone 2014). Microsatellites have also been utilized as markers of cells within organisms and tissues to reconstruct cell lineage histories (Reizel et al. 2012; Evrony et al. 2015; Wei and Zhang 2020).

The repetitive structure of microsatellites that makes them highly mutable *in vivo* and useful as lineage markers also has a downside: microsatellites incur a higher frequency of *in vitro* artifacts than other genomic elements, which can confound accurate genotyping (Selkoe and Toonen 2006). Microsatellites mutate by polymerase slippage during replication when transient denaturation of the replicating strand followed by incorrect reannealing to the wrong repeat leads to insertion or deletion of a repeat unit in the newly synthesized strand (Schlötterer 2000; Ellegren 2004; Bhargava and Fuentes 2010). This same process occurs during *in vitro* amplification of microsatellites, for example in PCR. This produces an artifact pattern called “stutter” in which the final population of amplified molecules has a distribution of repeat unit counts with a peak at the true repeat unit count (i.e., the true genotype). Amplified microsatellites can be accurately analyzed by capillary electrophoresis despite this “stutter” pattern, since the highest signal peak representing the true genotype can be readily distinguished from adjacent “stutter” peaks that have lower signal (Wenz et al. 1998; Acquaviva et al. 2003; R. Vemireddy et al. 2007). However, the throughput of capillary electrophoresis limits its use to small numbers of loci and samples (Hill et al. 2009; Guichoux et al. 2011).

High-throughput sequencing can profile many more microsatellites than capillary electrophoresis, however the sequencing must have sufficient read depth, generally higher than that required for calling base substitutions, for the true genotype to be apparent among the stutter distribution caused by library preparation artifacts and sequencing errors (Willems et al. 2017). This requirement for higher than standard sequencing read depth is further exacerbated when using prevalent short-read sequencers, since only a fraction of reads will fully span each microsatellite to enable genotyping of its number of repeats. The degree of microsatellite stutter artifact and the consequent read depth required for accurate genotyping can be mitigated by PCR-free sequencing (Fungtammasan et al. 2015), however, in many applications, the amount of DNA available is low and requires PCR amplification. While high-depth whole-genome sequencing (WGS) of PCR-amplified libraries could achieve large-scale, high-fidelity profiling of microsatellites, this is not feasible to scale to hundreds of samples per condition or experiment. Here, we developed an integrated suite of computational and novel molecular methods for targeted sequencing and analysis of > 150,000 microsatellite loci — achieving both cost-effective large-scale profiling of microsatellites and high fidelity (**Fig. 1A**). These tools may enable new applications of microsatellite profiling to a broad range of biological questions.

**Figure 1.**
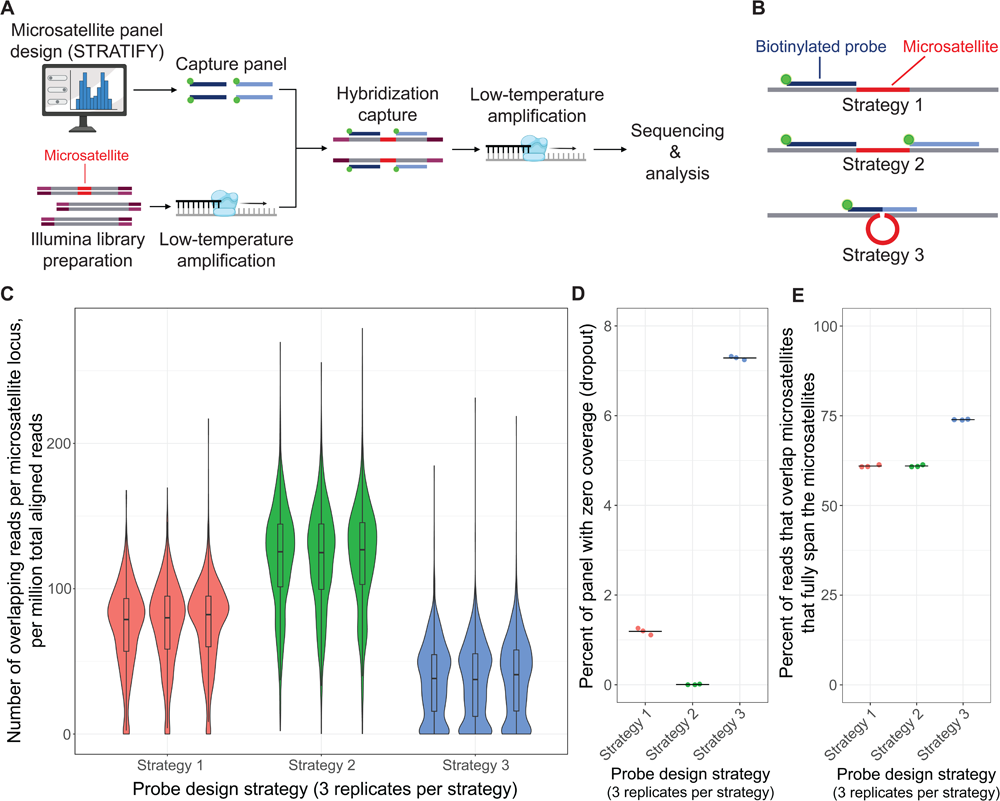
Overview of method and probe design strategy. **A.** Schematic of microsatellite panel design, library preparation, hybridization capture, sequencing, and analysis. **B.** The three probe design strategies tested in the pilot capture panel. **B.** Distribution of coverage across targeted microsatellite loci for each probe design strategy. Normalized coverage is plotted as the [number of microsatellite-spanning reads at the locus] / [number of million aligned reads in the sample]. Box-and-whiskers show the first quartile, median, and third quartile of the distributions. Whiskers show 1.5 x the interquartile range. **D.** Fraction of loci in the panel with zero coverage (i.e., “dropout”) for each probe design strategy. Mean value is represented by a black line. **E.** Percent of reads that overlap microsatellites that also fully span the microsatellites, for each probe design strategy. Calculated as [total number of reads fully spanning the targeted microsatellites] / [total number of reads overlapping targeted microsatellites by at least 1 bp] * 100. Mean value is represented by a black line. Note, **Figs. 1C-E** show experimental samples profiled using standard library amplification temperature.

## RESULTS

### Selection of microsatellite loci for profiling

We considered several factors when choosing microsatellite loci for targeted profiling. First, targeted profiling of microsatellites would benefit from capturing a panel of loci that are more likely to have mutations, because this would increase the power of resolving differences and lineage relationships among samples. Some microsatellite loci are less likely to be informative, because their mutation rates are relatively lower, such as loci with longer repeat motifs or lower repeat unit counts (Lai and Sun 2003; Ellegren 2004). Additionally, microsatellites must be in regions of the genome accessible to short reads, i.e., with good mappability. Microsatellites in low-mappability regions, specifically loci with low-mappability (non-unique) flanking sequences, would yield sequencing reads that cannot be aligned well to the genome with short reads, thereby precluding analysis. Furthermore, loci with low-mappability are poor targets for targeted profiling because the oligonucleotide probes used to target them would also hybridize to other undesired loci. For these reasons, it is essential that for any given microsatellite panel size, loci are selected that are most likely to be capturable, alignable to the genome, and informative. While many computational tools exist to identify microsatellite loci across the genome (Lower et al. 2018), these lack the necessary informatic annotations and the ability to rapidly iterate across many analytic parameters to select such an optimized panel of microsatellite loci. We therefore developed a user-friendly, web-based app called STRATIFY (Short Tandem Repeat Analysis and Target Identification) that enabled us to design an optimized, large-scale panel of human microsatellites for targeted profiling.

First, we used the Tandem Repeats Finder (TRF) tool (Benson 1999) to annotate microsatellites in the human genome, and we optimized TRF’s settings to maximize the number of annotated loci without introducing a large number of loci with highly imperfect motif repeats (**Figs. S1A-C** and **Methods**). We then annotated additional features of these microsatellite loci, including: a) uninterrupted length (i.e., the length, in bp, of the longest span of perfect repeats of the microsatellite base motif without any indels or mismatches) and uninterrupted copy number (i.e., the maximum number of consecutive perfect repeats of the base motif without any indels or mismatches), as these parameters have been shown to contribute to the microsatellite mutation rate (Sun et al. 2012; Willems et al. 2016); b) mappability (Karimzadeh et al. 2018) and GC content of the microsatellite and each of its 5’ and 3’ flanking sequences; c) overlap between a given microsatellite and its neighboring microsatellite or the distance to the nearest microsatellite; d) overlap with regions of the genome prone to alignment errors, for example segmental duplications (Bailey et al. 2002) and the ENCODE Data Analysis Center Blacklist (Amemiya et al. 2019); e) replication timing (Ryba et al. 2011; Marchal et al. 2018); and f) estimated mutation rate based on a model taking into account the uninterrupted length and motif size of any given microsatellite (Gymrek et al. 2017).

We next built an interactive web application, STRATIFY, that allows filtering of the annotated microsatellites based on any of TRF’s and our supplemental annotations (**Figs. S1D** and **S2**). STRATIFY also dynamically updates plots of data for visualization and quality control (**Fig. S1E**). This application was essential in allowing us to rapidly explore many combinations of parameters to select an optimal set of microsatellite loci for targeted profiling (**Methods**).

### Probe design for hybridization capture of microsatellites

We chose a hybridization capture approach for targeted profiling of microsatellites, because it is scalable to hundreds of thousands of loci as evidenced by its use in whole-exome sequencing (Majewski et al. 2011). This contrasts with other amplicon-based targeted sequencing approaches, such as molecular inversion probes, that are typically limited to < 20,000 loci (Mamanova et al. 2010), with the largest microsatellite panel to date profiling ∼25,000 loci (Campbell et al. 2015; Wei and Zhang 2020; Tao et al. 2021). Scaling beyond the limits of amplicon-based approaches is important for microsatellite profiling, since the power to resolve relationships among samples or cells increases with the number of mutations and consequently with the number of microsatellite loci profiled (Gärke et al. 2012).

Importantly, hybridization capture of microsatellites cannot be performed efficiently with probes (oligonucleotides complementary to target sequences) directly targeting the microsatellite repeat sequences. First, the sequence of a microsatellite is often imperfect, with varying base substitutions and indels across loci and samples that would affect probe affinity. Second, probes designed to microsatellite motifs themselves cannot be targeted to specific loci and would therefore capture all loci with that motif, including those with non-ideal features (e.g., low mappability). Additionally, the vast majority (97.6%) of microsatellites are shorter than the length of probes required by the temperature and stringency used to achieve specific genome-wide hybridization capture (55 bp – 120 bp) (Kruglyak et al. 1998; Samorodnitsky et al. 2015). Therefore, probes that span both the microsatellite sequence and extend into its flanks would have variable mismatches due to variability in microsatellite length and sequence. For these reasons, we developed a different approach for capturing microsatellites that does not target the microsatellite sequence itself.

The sequences of the flanks adjacent to a microsatellite are much less likely to be variable across samples than the microsatellite itself, and they are also specific to each locus. We therefore decided to target our hybridization capture probes to the flanks of microsatellites. However, since standard capture probes are designed to directly overlap their target sequences, we first tested whether probes designed to microsatellite flanks would adequately capture reads fully spanning the adjacent microsatellite to enable genotyping. Specifically, we developed and evaluated three different probe design strategies for microsatellite flanks (**Fig. 1B**). In strategy 1, one flank of the microsatellite was targeted by a single probe. In strategy 2, both flanks of the microsatellite were targeted, each by a separate probe. In strategy 3, both flanks were targeted using a single probe — the left and right halves of the probe targeted the left and right flanks, respectively. In strategy 3, each flank binds to the probe, and the microsatellite sequence remains unhybridized as a single-stranded DNA “loop”. The potential advantage, if successful, of this unconventional probe strategy compared to strategies 1 and 2 would be a higher probability of capturing DNA fragments that fully span the microsatellite, which is required for genotyping. We subsequently used STRATIFY to obtain a set of loci that passed our design filters and that could be captured with all three design strategies (**Fig. S3A**, **Table S1,** and **Methods**). We designed this first pilot panel to target only microsatellites with 2 to 4 bp motif lengths, because these loci are abundant and more mutable than those with longer motif lengths (Lai and Sun 2003). For each of the three strategies, we targeted 3,333 random loci from this set of loci.

We performed hybridization capture with this pilot panel on three replicates of control genomic DNA (Sample NA12877) (**Table S2**) and evaluated the performance of each probe design strategy. As an initial metric of capture quality, we evaluated the distribution of coverage by reads fully spanning each microsatellite locus (**Fig. 1C**). Strategy 2 (both flanks targeted, each by a separate probe) had a significantly higher distribution of coverage than strategy 1 (one probe targeting one flank) and strategy 3 (one probe targeting both flanks), and strategy 3 had the lowest distribution of coverage. In strategy 2, only 0.005% of loci on average had complete dropout (zero coverage), while strategies 1 and 3, had 1.2% and 7.3% complete dropout, respectively (**Figs. 1D** and **S5A**). The distribution across loci of the ratio of microsatellite coverage to flank coverage was also significantly higher for strategy 2 than strategy 1 (**Fig. S4**), consistent with targeting of both flanks more efficiently capturing DNA fragments containing the microsatellite. Notably, this ratio was highest for strategy 3, suggesting that when a locus is captured well by this design approach, a greater proportion of captured DNA fragments contain the microsatellite (**Fig. S4**). However, this positive metric for strategy 3 is tempered by this design strategy’s significantly worse absolute coverage (**Figs. 1C-D**), indicating that its absolute capture efficiency is lower than that of the other probe design strategies. Strategy 2 also performed best in terms of AT dropout, a measure of how under-covered AT-rich regions are compared to expectations based on the reference genome (Zhou et al. 2021), and fold 80 base penalty, a measure of capture uniformity (**Figs. S5B-D**).

Overall, these analyses indicate that strategy 2, in which both flanks are targeted each by a separate probe, achieves the best capture performance for microsatellites, with nearly all loci captured with high average coverage. While strategy 1 also performed well, especially in sequencing uniformity, its overall performance was lower than Strategy 2. Strategy 3 had the worst performance, with higher dropout and lower uniformity; while it achieved our goal of increasing the proportion of reads fully spanning the microsatellite (74% of reads overlapping targeted microsatellites by at least 1 bp also fully spanned the microsatellites, compared to 61% for both strategies 1 and 2 [**Fig. 1E**]), this was counteracted by decreased overall capture efficiency (**Figs. 1C-D** and **S5A-D**). We therefore chose to proceed with strategy 2 whenever a locus allowed design of probes to both flanks (based on design filters) and strategy 1 for loci that have only one flank passing our design filters. This combined approach using strategies 1 and 2 significantly increases the number of loci available for capture, because only one flank needs to pass our filters (**Fig. S3B**) and most loci are still captured efficiently with only one probe (**Fig. 1C-D** and **S5A-D**).

### Low-temperature amplification reduces microsatellite stutter artifacts

One of the most challenging aspects of microsatellite sequencing is the production of stutter artifacts during *in vitro* amplification (Ellegren 2004; Fungtammasan et al. 2015). *In vivo*, errors in microsatellite replication are corrected by mismatch repair machinery (Ellegren 2004), which is absent from *in vitro* amplification. *In vitro* amplification-induced stutter can obscure the true genotype of a microsatellite locus and confound downstream analyses. While targeted hybridization capture can significantly improve the scalability of profiling microsatellites, it creates the additional challenge of requiring two *in vitro* PCR amplifications, before and after the hybridization capture. Therefore, we explored methods for reducing stutter artifact during *in vitro* amplification that could be applied to genomic libraries.

Stutter events that occur in early PCR cycles contribute more to the final stutter distribution. To mitigate this issue, we performed linear amplification cycles at the beginning of the PCR reaction that would prevent exponential expansion of stutter events. However, solely using linear amplification in our PCR protocol would not provide enough product for downstream steps of the protocol. We therefore decided to begin our protocol with 3 cycles of linear amplification, then add the second primer to initiate exponential PCR for the remainder of the reaction. To reduce the possibility of stutter events during this phase, we optimized our protocol to use the fewest possible number of amplification cycles required to achieve our desired DNA yield. We were able to reduce our cycle count to only 3 linear and 2 exponential cycles for the pre-hybridization PCR, and 3 linear and 8 exponential cycles for the post-hybridization PCR (**Fig. 2A** and **Methods**).

**Figure 2.**
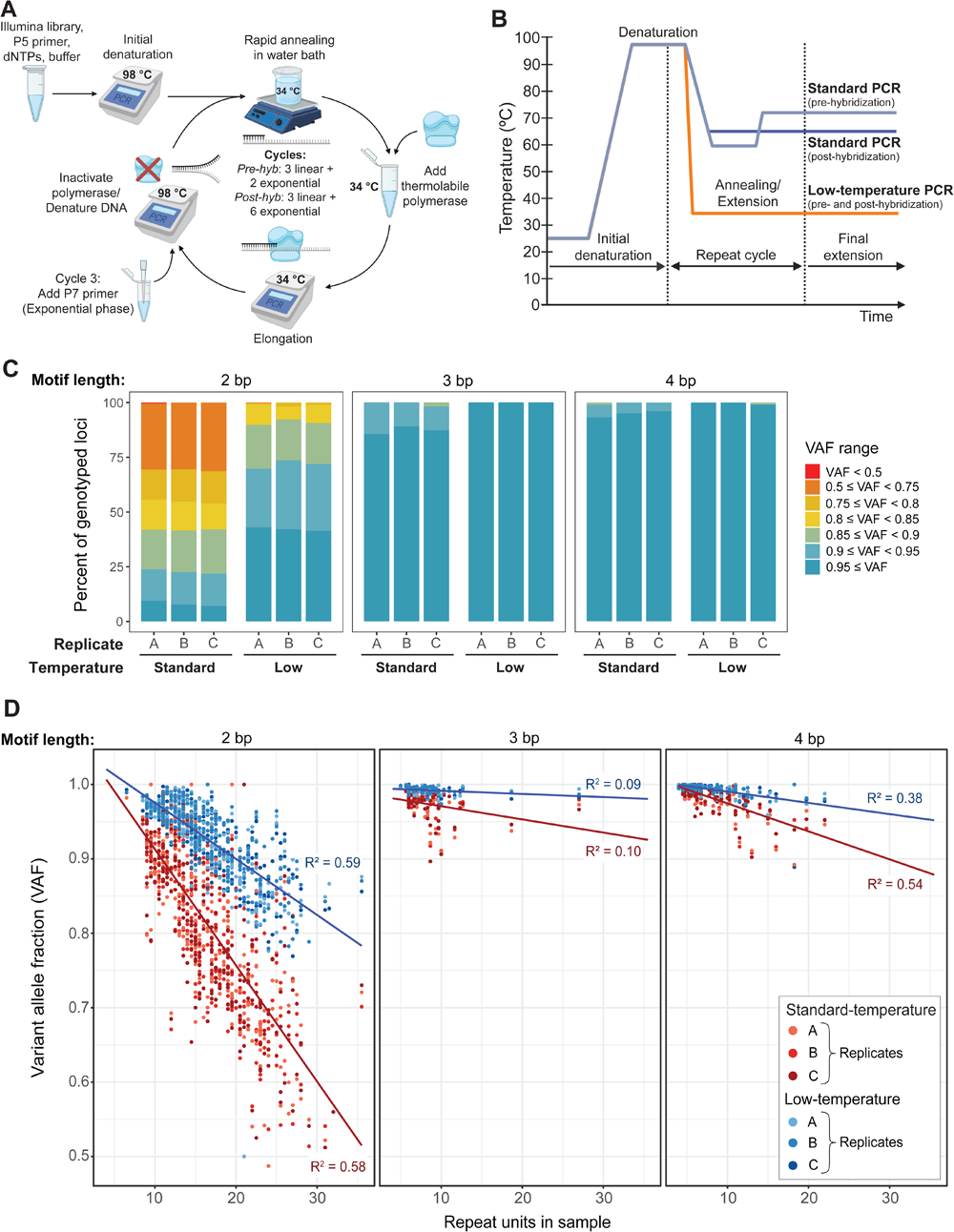
Impact of amplification temperature on microsatellite stutter. **A.** Schematic of low-temperature PCR of sequencing libraries that significantly reduces microsatellite stutter artifact. Pre-hyb, pre-hybridization PCR; post-hyb, post-hybridization PCR. The number of cycles specified is for the large-scale capture panel (**Methods**). **B.** Schematic graph of temperatures during standard and low-temperature PCR protocols. Pre-hybridization (pre-hyb) and post-hybridization (post-hyb) standard PCRs use different thermal cycling protocols. In both standard and low temperature PCR, the pre-hybridization protocol consisted of 3 linear and 2 exponential PCR cycles, and the post-hybridization protocol consisted of 3 linear and 5 exponential PCR cycles. **C.** Fraction of genotyped loci on male chromosome X microsatellite loci in 3 replicates of sample NA12877 (male) in different bins of variant allele fraction (VAF). Only chromosome X loci from male individuals are included in the analysis to provide accurate stutter estimates, which is not feasible for bi-allelic loci. Loci are grouped by motif length, because motif length correlates with mutability. VAF = [number of reads supporting the allele genotyped by HipSTR] / [total number of reads at the locus calculated as the sum of the HipSTR MALLREADS field]. A VAF < 1 indicates stutter reads at the locus. Low-temperature amplification significantly decreased the fraction of loci with high levels of stutter (i.e., with low VAF), most noticeably for 2 bp motifs. **D.** VAF of genotyped chromosome X loci (as calculated in Fig. 2C) versus the number of repeat units called in the sample in 3 standard-temperature (red) and 3 low-temperature (blue) amplification replicates of individual NA12877. Red line: linear regression for VAF values in the standard-temperature condition. Blue line: linear regression for VAF values in the low-temperature condition.

We also suspected that temperature was one of the most important factors for reducing the frequency of stutter events, based on two prior studies that observed significantly decreased stutter when using low-temperature or isothermal amplification protocols (Hite et al. 1996; Daunay et al. 2019). In standard PCR, extension is typically conducted between 55 °C and 72 °C. At these elevated temperatures, the synthesized and template strands are more likely to denature and “slip” during reannealing, causing insertion or deletion of repeat units (i.e., stutter). If the temperature is lowered during extension, the microsatellite strands may be less likely to denature and cause stutter events. Hite et al. (Hite et al. 1996) applied low-temperature amplification to one microsatellite locus by adding a thermolabile polymerase at each cycle with extension at 37 °C, and they observed reduced stutter bands in gel electrophoresis compared to amplification at high temperature with thermostable polymerase. Daunay et al. (Daunay et al. 2019) applied recombinase polymerase amplification (RPA), an isothermal amplification method, and observed a similar reduction in stutter by capillary electrophoresis and amplicon sequencing, even when the reaction was multiplexed for three loci. However, low temperature amplification has not yet been applied to large-scale microsatellite genomic libraries.

Here, we further developed the above low-temperature PCR method for large-scale microsatellite libraries (**Figs. 2A-B**). In our method, a thermolabile polymerase is added at each cycle of amplification, with extension occurring at 34 °C. Denaturation in each cycle occurs at 98 °C followed by rapid cooling in a water bath to minimize extension by any residual polymerase activity that survives the heat denaturation (Hite et al. 1996). The combination of this low-temperature annealing and extension, linear amplification, and fewer amplification cycles should significantly lower the amount of stutter observed in genomic sequencing libraries.

To test the ability of our low-temperature amplification method to reduce stutter in a genomic library, we profiled three technical replicates of NA12877 with the above pilot panel using low-temperature for both the pre-and post-hybridization capture amplification steps. Note that the standard-temperature pilot panel samples of the prior section and these low-temperature pilot panel samples were profiled identically, except for temperature, in the same experiment on the same day. We then compared the stutter levels of standard-temperature and low-temperature samples for loci on chromosome X, because in male samples like NA12877, chromosome X loci are single-allele loci and any read not matching the genotype reflects stutter. By contrast, stutter cannot be readily measured in biallelic loci (i.e., autosomal loci), because the stutter distribution of one allele may overlap the true peak of the other allele.

We found that low-temperature amplification significantly reduced the fraction of loci with high levels of stutter (**Fig. 2C** and **Table S3**), as measured by the fraction of reads supporting the called genotypes (variant allele fraction, VAF; higher VAF = lower stutter). Across all technical replicate of both temperature conditions, an average of 432 (range: 429 – 437) chromosome X loci were genotyped. In the low-temperature samples, an average of 0.08% of genotyped loci had a VAF less than 0.75, while in the standard-temperature samples, an average of 19.6% of genotyped loci had a VAF less than 0.75. All loci with a VAF less than 0.75 had a motif length of 2 bp. In both standard-and low-temperature conditions, all 3 and 4 bp loci had VAFs of 0.85 or higher, although the low-temperature samples still had less overall stutter than the standard-temperature samples (**Table S3**). Of note, 93% of stutter reads were shorter than the called allele (**Table S3**), consistent with studies showing that PCR stutter preferentially decreases rather than increases microsatellite length (Ellegren 2004). Furthermore, the increased stutter seen in loci with a 2 bp motif length is consistent with prior studies that found a higher probability of microsatellite mutation for shorter motif lengths (Kruglyak et al. 1998; Ellegren 2004; Bhargava and Fuentes 2010; Fungtammasan et al. 2015).

We also observed that for each motif length, low-temperature amplification achieved lower stutter levels than standard-temperature amplification across all total microsatellite lengths (**Fig. 2D**). Within each motif length, longer microsatellites (i.e., loci with more repeat units) also generally had higher stutter levels — again consistent with prior studies (Ellegren 2004; Bhargava and Fuentes 2010; Sun et al. 2012) (**Fig. 2D**). Altogether, these findings show that temperature is the most important factor identified to date in reducing stutter artifact during microsatellite amplification. Additionally, there were no significant differences in the distribution of coverage across loci or the quality of hybridization capture between standard-and low-temperature amplification (**Figs. S5A-D**). Overall, our low-temperature amplification method achieves, for the first time, a significant reduction in stutter artifacts for all types of microsatellites in highly multiplexed, large-scale genome sequencing libraries. This represents an important step towards large-scale, accurate genotyping of microsatellites.

### Large-scale profiling of microsatellites

After developing an optimal probe design strategy and a low-temperature amplification method for reducing stutter with a small-scale pilot panel of loci, we next proceeded to scale our method to a much larger number of microsatellite loci. A larger number of loci in the capture panel would better resolve differences among individuals or cells by increasing the number of detected mutations. Using our optimized approach for probe design (targeting both microsatellite flanks when possible, and only one flank when not possible) and an optimized set of filters, we produced a large-scale panel of 154,188 microsatellite loci (**Fig. 3A** and **Table S4**). Similar to the pilot panel, this panel included microsatellites with 2 to 4 bp motif lengths (147,656 loci) because of their relatively increased mutability compared to penta-and hexa-nucleotide loci (Lai and Sun 2003). The panel also included 6,532 poly-A loci, which have even higher mutation rates than loci with 2-4 bp motifs (Ellegren 2004). Because poly-A loci incur more significant stutter artifacts that make them challenging to genotype, especially in diploid chromosomes where the signals of two alleles need to be deconvoluted, we only included poly-A loci on chromosome X (**Fig. S6A**). We reasoned these may be amenable to genotype at least in male samples containing only a single chromosome X. At the same time, our low-temperature amplification method with reduced stutter motivated the possibility that poly-A loci may be feasible to genotype even in females. The size of this panel (28 Mb total probe length) is on the order of a whole-exome capture panel (∼ 40 Mb) (Zhou et al. 2021), making it the largest scale targeted profiling of microsatellites to date.

**Figure 3.**
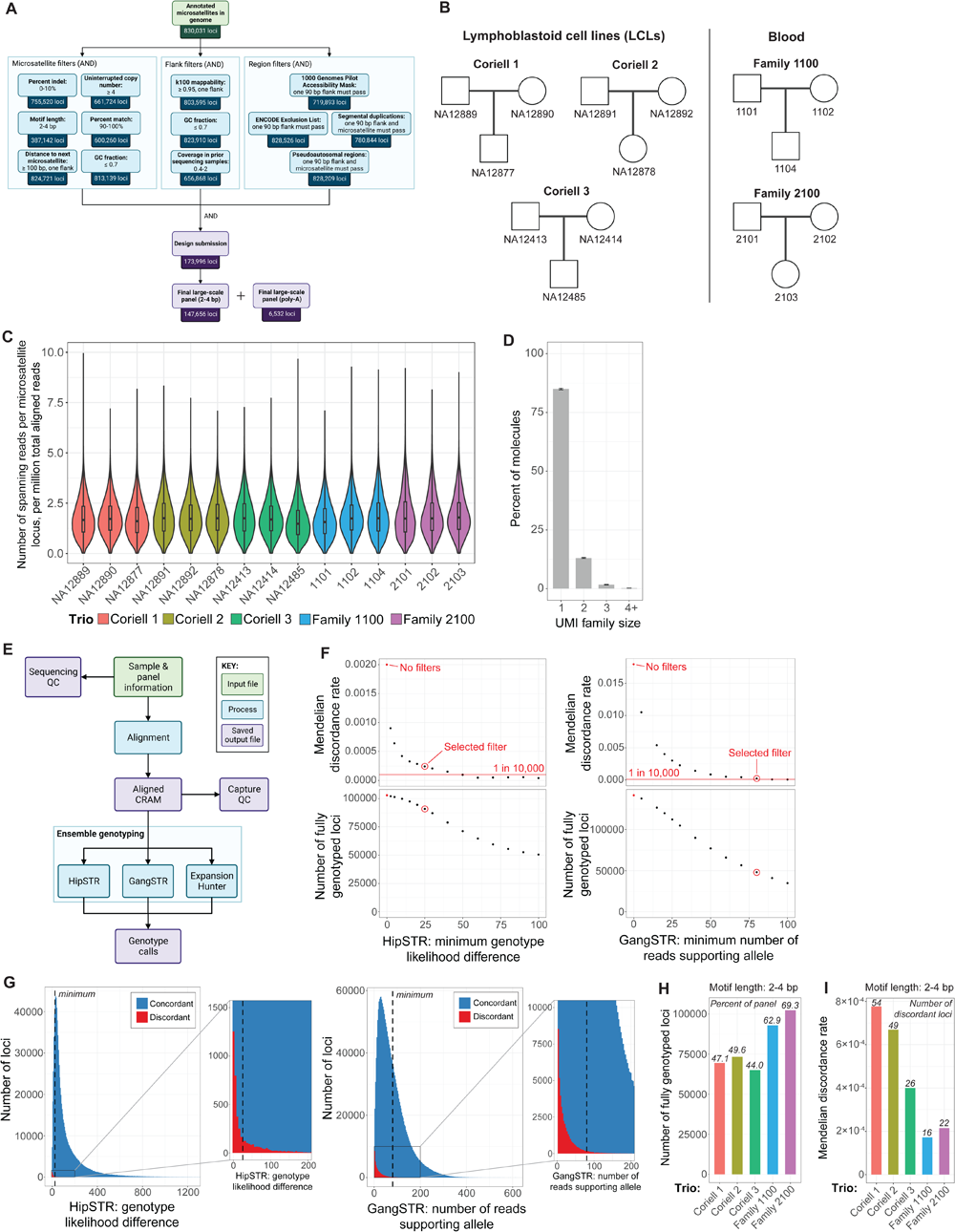
Large-scale microsatellite capture, genotyping pipeline STREAM, and Mendelian discordance analysis. **A.** Schematic of selection of loci with 2 to 4 bp motif lengths for the large-scale microsatellite capture panel. All filters are applied with Boolean AND logic. Different filtering parameters were used for poly-A loci in the panel (**Fig. S6A**). QC, quality control. **B.** Five family trios profiled with our large-scale microsatellite panel. **C.** Distribution of coverage across targeted microsatellite loci for the large-scale panel. Trios are arranged by trio in the following order: father, mother, child. Normalized coverage is plotted as the [number of microsatellite-spanning reads at the locus] / [number of million aligned reads in the sample]. Box-and-whisker plot shows first quartile, median, and third quartile of distribution. Whiskers show 1.5 x the interquartile range. **D.** Average unique molecular identifier (UMI) family size (number of reads containing the same UMI) for libraries from 6 samples (Family 1100 and Family 2100) for which we also sequenced UMIs. A UMI family size of 1 indicates that the molecule was sequenced only once; family size of ≥ 2 indicates multiple sequencing reads for the same original input DNA molecule. Error bars: 2 standard deviations from the mean. **E.** Simplified schematic of STREAM, a microsatellite ensemble genotyping pipeline that optimally integrates calls from HipSTR, GangSTR, and ExpansionHunter. A detailed schematic of the pipeline is shown in **Figs. S6C-D**. **F.** Representative plots of filter parameter optimizations for STREAM based on Mendelian discordance rates and the number of loci genotyped at different thresholds. Data is shown for Family 1100. Red dot: values without quality filtering. Red circle: threshold value chosen for final filtering settings. Red line: approximate expected microsatellite de novo mutation rate (i.e., Mendelian discordance rate) based on prior studies (Weber and Wong 1993; Huang et al. 2002; Sun et al. 2012; Kristmundsdottir et al. 2023). Note that individual parameter filters do not achieve the expected discordance rate, and the final filter settings utilize an optimized combination of filters to achieve the expected discordance rate (**Methods**). See **Fig. S9** for plots of other filter parameter optimizations. **G.** Overlayed histograms of parameter values across all loci for two of the filtering parameters we optimized. Histograms are generated from data from all 15 samples captured with the large-scale panel. Plots include only fully genotyped loci, as only these loci have concordance calls. Concordant calls are blue and discordant calls are red. The zoom plot shows the distribution of discordant calls near the final filtering threshold. Dashed line: chosen threshold in our final filter settings. See **Fig. S10** for similar plots of other filter optimization parameters. **H.** Number of fully genotyped 2 to 4 bp motif length loci for each family trio; i.e. loci with genotypes called for all three members of the trio. Italics: fraction of capture panel that is fully genotyped. **I.** Mendelian discordance rate of fully genotyped loci with 2 to 4 bp motif lengths in each family trio. Mendelian discordance rate = [number of discordant loci] / [number of fully genotyped loci]. Italics: number of discordant 2 to 4 bp motif length loci.

We applied this large-scale panel to five family trios (father, mother, child; **Fig. 3B** and **Table S5**), since Mendelian discordance analyses can be used for high-fidelity assessment of our method’s accuracy as well as for optimization of analytic parameters. DNA from three trios was derived from lymphoblastoid cell lines (LCLs), and DNA from the other two trios was derived from blood samples. Either 3 or 6 samples were multiplexed per hybridization capture, and we sequenced the resulting libraries to an average of 873 total reads per locus, and an average of 218 reads fully spanning the microsatellite per locus. The distribution of coverage across loci was consistent across samples, indicating that our capture panel and low-temperature amplification are reliable and reproducible methods even at very large scale (**Fig. 3C**). An average of 93.6% of reads were on-target (i.e., aligned to the targeted microsatellites), and dropout was very low (average of 1.7% of loci with zero coverage) (**Figs. S7A-B** and **Table S5**). AT dropout averaged 8.1%, similar to the pilot panels for probe design strategies 1 and 2 (**Figs. S5C** and **S7C**). Finally, the average fold 80 base penalty was 1.6, which was similar to the fold 80 base penalty for strategy 1 in the pilot panel (**Figs. S5D** and **S7D**).

We also assessed the level of molecular duplication in the sequencing data. Duplicate sequencing reads of the same original input DNA molecule may differ due to stutter artifacts. If molecular duplication levels were very high (e.g., > 20 duplicates per molecule), this could enable creation of a high-fidelity consensus sequence for each original DNA molecule. However, achieving this level of duplication per molecule would require more than an order of magnitude more sequencing per sample, which is not scalable. In the absence of this, ideally there would be minimal duplication of molecules so that sequencing reads provide independent observations of a larger number of input DNA molecules. To assess the level of molecular duplication, we sequenced unique molecular identifiers (UMIs) that are present in our library adapters and found that nearly all molecules (98.1%) had only 1 or 2 molecular copies in the final sequencing data (**Figs. 3D** and **Fig. S6B**). This very low molecular duplication rate indicates that our capture is highly efficient and that our optimized low number of PCR cycles maintains high library complexity. It also suggests that in the future, it may be feasible to multiplex more than six samples per hybridization capture for even better cost efficacy.

As a final evaluation of our large-scale panel’s performance in terms of stutter artifact, we analyzed, for each motif length, the fraction of loci genotyped in different VAF ranges (**Fig. S8** (per above, higher VAF indicates lower stutter). As for the pilot panel, this analysis examined only chromosome X loci (8 of the trio individuals were male) to obtain accurate estimates of stutter. For 2 to 4 bp loci, we saw similar stutter levels as in the pilot panel, confirming that the size of the panel does not impact the ability of our low-temperature amplification method to reduce stutter (**Fig. S8**). Similar to the pilot panel, the fraction of loci with low VAF values decreased with increasing motif length (**Fig. S8**). Poly-A loci had more stutter than 2-4 bp motif loci, but most genotyped poly-A loci (84.6%) still had a VAF ≥ 0.75 (**Fig. S8**).

### Ensemble computational pipeline for genotyping microsatellites

While we achieve efficient capture of > 150,000 microsatellite loci with low levels of stutter, calling genotypes from microsatellite sequencing data is challenging and requires specialized approaches (Gymrek et al. 2012; Treangen and Salzberg 2012). Indeed, microsatellite stutter artifacts and the increased error in calling insertions and deletions in sequencing data have led several groups to develop tools specifically for genotyping microsatellites (Treangen and Salzberg 2012; Fang et al. 2014; Raz et al. 2019). Preliminary application of a few of these tools — HipSTR (Willems et al. 2017), GangSTR (Mousavi et al. 2019), and ExpansionHunter (Dolzhenko et al. 2019) — to our data revealed complementary strengths and weaknesses. For example, while HipSTR produces accurate calls as seen in prior studies (Halman and Oshlack 2020; Oketch et al. 2023), it does not output a genotype for many loci (22% of loci on average in our samples) due to HipSTR being unable to perform local realignment of the flanks. On the other hand, GangSTR and ExpansionHunter can call genotypes of loci that HipSTR does not call, but their accuracy is lower, especially at lower read depths (Halman and Oshlack 2020; Oketch et al. 2023). We therefore developed a computational pipeline, STREAM (Short Tandem Repeat Ensemble Analysis Method; schematic in **Fig. 3E** and detailed schematic in **Figs. S6C-D**), that combines calls from all three of these tools to genotype as many loci as possible while maintaining high call quality. Additionally, our trio experimental design provided a very low-level true signal of de novo mutations (discordant child-parent calls) that allowed us to optimize our pipeline’s parameters based on expected de novo mutation rates from prior trio studies, which was then followed by our own experimental validation of candidate de novo mutations called by our pipeline.

STREAM first aligns reads and then runs HipSTR, GangSTR, and ExpansionHunter. We run HipSTR jointly across all samples since this allows generation of more accurate stutter models that improve accuracy (Willems et al. 2017), while GangSTR and ExpansionHunter can only process each sample individually. Next, the genotypes from all three callers are collated. Since each of the three callers may have biases in genotyping at each locus, we choose for each locus one caller for all samples. The algorithm for deciding which caller to use for each locus was based on systematic optimization of filter parameter values and combinations that was guided by both the expected Mendelian discordance rates from prior studies and the number of loci that were “fully genotyped” (i.e., genotyped in all members of the trio) (**Methods**). We similarly optimized the order of which callers to prioritize in the ensemble genotyping process. Specifically, the filter optimization process identified a set of parameters that produced the expected Mendelian discordance rate based on prior estimates of microsatellite de novo mutation rates (∼ 1 in 10,000 loci) (Weber and Wong 1993; Huang et al. 2002; Sun et al. 2012; Kristmundsdottir et al. 2023) and that maximized the number of “fully genotyped” loci. We used Family 1100 for this initial optimization because it was a trio of blood samples, and we expected blood samples to have a lower discordance rate than LCLs due to the possibility of microsatellite mutations occurring during cell culture. Family 1100 was also sequenced more deeply than the LCL trios and more evenly across all three trio members than Family 2100.

We first explored a range of threshold values independently for each filter to assess how this affected the rate of child-parent discordant calls and the number of fully genotyped loci (**Fig. 3F** and **Figs. S9A-J**). Ideally, a filter would remove many more discordant calls than concordant calls (when above the expected discordance rate due to de novo mutations) without filtering out a large fraction of loci. After identifying parameters that help lower the discordance rate, such as mean quality, the difference in genotype likelihood between the called genotype and the next most likely genotype, and the number of reads supporting the called alleles, we tested different combinations of these filters and caller preference orders until we achieved the expected low-level Mendelian discordance rate while maximizing the number of “fully genotyped” loci. Subsequently, we plotted the distribution of concordant and discordant calls across all trios to fine-tune the final filtering thresholds (**Fig. 3G** and **Fig. S10A-I**). This filtering was performed separately for loci with 2 to 4 bp motif lengths and poly-A loci because the higher stutter of poly-A loci requires more stringent filtering thresholds. Additionally, for loci with 2 to 4 bp motif lengths, we monitored the balance of each motif length in the final set of genotypes versus the full panel to ensure that our filtering does not significantly bias the genotyped motif lengths (**Fig. S11A**).

Our final optimized pipeline produced an average of 55% and 57% fully genotyped loci for 2 to 4 bp motif loci and poly-A loci, respectively (**Fig. 3H** and **S11B**). Three trios (Coriell 1, Coriell 2, and Coriell 3) had fewer fully genotyped loci (47% and 51% on average for 2-4 bp motif and poly-A loci, respectively) compared to the other two trios (66% and 66% on average, respectively; Family 1100 and Family 2100) due to the former trios’ average 31% lower sequencing depth (**Table S5**). This suggests that higher read depth increases the number of fully genotyped loci.

Importantly, for loci with 2 to 4 bp motif lengths, STREAM measured an average Mendelian discordance rate of 1.93 × 10^-4^ for trios whose DNA derived from blood (range: 1.72 × 10^-4^ – 2.15 × 10^-4^) and 6.15 × 10^-4^ for the trios whose DNA derived from LCLs (range: 4.00 × 10^-4^ – 7.77 × 10^-4^) (**Fig. 3I**). These results are consistent with expected microsatellite de novo mutation rates (Weber and Wong 1993; Huang et al. 2002; Sun et al. 2012) and the accumulation of additional mutations during divergence of the trio LCLs in culture. Loci with a motif length of 2 bp had the highest discordance rate in all trios, consistent with the higher mutability of 2 bp motifs compared to 3 and 4 bp motifs (Ellegren 2004) (**Fig. S11D**). Additionally, as expected for the more mutable poly-A loci, we found they had an 8.1-fold higher discordance rate on average relative to 2 to 4 bp motif loci across all trios (**Figs. 3I** and **S11C**). Overall, these results support STREAM’s ability to call microsatellites with high fidelity across a large panel of microsatellites.

To further demonstrate the specificity of STREAM, we validated discordant loci called in the trios Coriell 3, Family 1100, and Family 2100 using capillary electrophoresis (**Table S6**). Capillary electrophoresis validated 97% (59 of 61) of loci with 2 to 4 bp motifs that were called as Mendelian discordant by STREAM. Three additional 2 to 4 bp motif loci were excluded from the validation, because upon manual review of sequencing data, we found that these apparent Mendelian discordant calls were due to a heterozygous deletion in one of the parents that was inherited by the child. The two loci that did not validate by capillary electrophoresis were the result of miscalls by genotyping tools, since manual review of sequencing data was concordant with capillary electrophoresis. Poly-A loci are more difficult to validate by capillary electrophoresis, due to their broad stutter patterns. Nevertheless, we validated 12 of 13 poly-A loci called as Mendelian discordant by STREAM whose capillary electrophoresis results were interpretable (6 additional loci were excluded from analysis due to broad stutter patterns that were not interpretable). These validation results demonstrate the high specificity of our method to detect rare microsatellite mutations.

Since total read depth is an important component for cost-effective scaling of our method to many samples, we further evaluated the dependence of our method’s performance on read depth. We subsampled the data for each member of Family 1100 from 30X to 1013X average reads per locus (1013X is the average read depth of sample 1101, the lowest coverage sample from Family 1100). We found that the Mendelian discordance rate remained stable regardless of sequencing depth, but the number of fully genotyped loci was highly dependent on sequencing depth (**Fig. S12A**). At 200X average coverage per locus, only one-quarter of loci are fully genotyped compared to the almost two-thirds of loci that are fully genotyped at full read depth (1013X), which may explain why discordant loci are not observed below 200X coverage, as they may not have been genotyped or passed our quality filters. These results emphasize the importance of high sequencing depths to genotype most loci and further highlights the importance of our targeted approach that makes these sequencing depths feasible.

To evaluate the cost-effectiveness of our method compared to PCR-free whole genome sequencing, we also analyzed high read depth PCR-free WGS data from a family trio (Zook et al. 2016). We subsampled each family member’s data from 30X to 309X average coverage (309X is the average read depth of the child’s sample, the lowest coverage sample in the trio). We analyzed the trio using either the loci targeted by our large-scale capture panel or a larger genome-wide set of 563,770 microsatellite loci and calculated the cost per thousand fully genotyped loci (**Figs. S12A-B**, **Supplemental Note,** and **Methods**). When analyzing only the loci from our large-scale panel, our method was significantly more cost effective than WGS at all coverage levels necessary to achieve detection of Mendelian discordant loci. When analyzing the genome-wide set of microsatellite loci, WGS achieves a comparable cost per thousand fully genotyped loci (at a read depth required for reliable detection of Mendelian discordant loci) with an approximately 3.2-fold higher number of fully genotyped loci, albeit with a 3.8-fold higher cost per sample (**Fig. S12A**). Furthermore, our targeted capture method becomes increasingly more cost effective as average read depth increases, which may be needed for some applications to maximize the number of samples genotyped at each locus (**Figs. S12A-B**).

## Discussion

Microsatellites are the most mutable genomic elements, and their consequent variation has enabled diverse applications in population genetics, forensics, and lineage tracing. Here, we introduce four tools and innovations that will increase the scalability and resolution of microsatellite profiling: 1) STRATIFY — a tool for designing large-scale microsatellite capture panels; 2) a novel method for the largest-scale targeted capture of microsatellite loci, to date; 3) a method for low-temperature amplification of large-scale microsatellite libraries that significantly reduces stutter artifact; and, 4) STREAM — an ensemble genotyping pipeline integrating three microsatellite callers for more comprehensive and accurate genotyping.

Current microsatellite profiling methods are limited by their scalability. Capillary electrophoresis can only process a small number of loci, while WGS can profile microsatellites genome-wide, but the read depths required for accurate genotyping are not currently feasible to scale to hundreds of samples per experiment. One of the most important advantages of our method is its ability to readily scale to > 150,000 loci, comparable in size to a whole-exome panel, which is important for maximizing the number of captured mutations for meaningful biological analyses. Our method captures significantly more loci than other recently reported methods, the largest of which captures ∼12,000 loci per sample (Wei and Zhang 2020; Tao et al. 2021). Additionally, we can capture loci with diverse motifs from across the genome, in contrast to methods that focus on specific motifs (Wang et al. 2022) or that utilize restriction enzymes to fragment DNA, which limits the number of targetable loci (Wei and Zhang 2020).

Stutter has long been one of the most challenging problems in microsatellite studies. Prior methods have achieved lower stutter for small numbers of loci (Hite et al. 1996; Daunay et al. 2019) or for specific microsatellite motifs such as ‘G’ homopolymers (Wang et al. 2022). Notably, our low-temperature amplification method achieves, for the first time, significant reduction of stutter in a large, complex sequencing library across many types of microsatellite motifs and motif lengths.

Concomitant with our novel molecular methods, we introduce two analytic tools, STRATIFY and STREAM, that facilitate the design and analysis, respectively, of large-scale microsatellite panels. STRATIFY allows for rapid iteration of different parameter thresholds for facile curation of panels, while STREAM combines the strengths of three microsatellite genotyping tools while mitigating their individual weaknesses to call genotypes accurately. Together, these advances will not only enable scaling of current microsatellite-based applications, but may enable novel applications, for example high-resolution single-cell lineage tracing (Wei and Zhang 2020; Tao et al. 2021). During preparation of this manuscript, a new tool called EnsembleTR was published that also utilizes HipSTR, GangSTR, and ExpansionHunter for ensemble genotyping of microsatellites (Ziaei Jam et al. 2023). STREAM and EnsembleTR differ in their analysis process. STREAM utilizes more fine-grained optimization of filter settings for each individual caller and then chooses the final genotype per an optimized order of caller preference, whereas EnsembleTR performs lenient filtering for each individual caller and then chooses the final genotype based on quality scores summed across individual callers. EnsembleTR’s approach requires it to reconcile differences in allele representations of each locus across the individual callers, whereas STREAM selects the final genotype directly from individual callers. Despite these differences, both tools highlight the advantage of an ensemble approach for maximizing genotyping accuracy for microsatellites and the number of loci genotyped.

Homopolymers are often excluded from microsatellite profiling methods due to their significant stutter artifacts. Here, we were able to successfully capture and genotype poly-A homopolymer loci due to the lower level of stutter achieved by our low-temperature amplification. Our success in genotyping poly-A loci on chromosome X in females (in Family 2100, 51% of loci were fully genotyped with a Mendelian discordance rate of 6.1 × 10^-4^) suggests that we could extend profiling of poly-A loci to all chromosomes. Given that poly-A repeats are the most abundant and most mutable type of microsatellite in the genome, including more poly-A loci in future panels would increase the number of mutations observed per sample and further improve the resolution of analyses.

We analyzed the cost effectiveness of our method compared to PCR-free WGS and found that our method is more cost effective than WGS at high read depths, or when a smaller number of loci need to be .profiled for the biological application (**Figs. S12A-B** and **Supplemental Note**). Therefore, the most cost-effective method depends on two key considerations: a) the number of genomic loci required by the biological application, and b) the importance of maximizing the fraction of samples genotyped at each locus; for example, lineage tracing applications would benefit from a higher read depth that maximizes the number of samples genotyped at each locus. Samples with baseline stutter that require higher read depth for accurate genotyping, such as amplified single-cell genomes, may also benefit from the higher cost-effectiveness of our method at higher read depths. Additionally, for applications requiring fewer loci (e.g., molecular markers for ecology), our method is far more cost effective than WGS. At the same time, as sequencing costs continue to decrease, high-depth PCR-free WGS has benefits in terms of ease of sample preparation, uniformity of genomic coverage, and profiling of more microsatellites per sample, such that it is likely to become the preferred method for applications requiring profiling of hundreds of thousands of loci.

Though we were able to reduce the level of stutter in our final libraries, we still observe a low-level stutter signal that confounds some genotype calls, and our filtering to remove these false calls eliminates the corresponding loci from the final analysis. Further reduction of stutter through temperature modification is unlikely. A recent study reduced stutter by targeted mutagenesis (Wang et al. 2022), but is limited to ‘G’ homopolymer microsatellites that would not be compatible with the scale of capture we have achieved. Thus, stutter continues to be a challenge despite the improvements we have demonstrated, motivating further research to address it, perhaps by novel polymerases (Yamanoi et al. 2021) or accessory proteins during library amplification.

One limitation of our method is that the low-temperature amplification requires manual transfer of samples between temperature conditions and addition of enzyme each cycle. Automation of this amplification process is feasible, either via liquid-handling robots or with microfluidics (Ahrberg et al. 2016), and this would lower the barrier to implementing this protocol while also enabling higher throughput. Another limitation of our method is that STREAM, our ensemble microsatellite genotyping tool, is limited by the accuracy and genotyping capabilities of the three microsatellite callers it utilizes. For example, HipSTR does not genotype some loci due to failure in local realignment. Our open-source pipeline can be readily adapted to incorporate new microsatellite genotyping tools, while our trio sequencing datasets could then be used to refine these tools’ new parameters.

The ability of our method to profile > 150,000 microsatellites across many samples with high fidelity may help elucidate previously opaque relationships among individuals in a population or among cells using mutational processes that are ubiquitous in all organisms and cells. Capture of the high endogenous levels of microsatellite mutations across many loci will allow for reconstruction of these relationships at a higher resolution than would be achievable with other markers that mutate less frequently. Our method also opens new opportunities to study non-model organisms without the need for engineered molecular markers, which is especially important for retrospective studies of human tissues.

## METHODS

### Microsatellite panel design tool: STRATIFY

#### Identifying microsatellite loci in the genome

We identified microsatellite loci in the hg38 human reference genome with Tandem Repeats Finder (TRF) version 4.09.1 (Benson 1999). We selected TRF because it provided the greatest flexibility in tuning microsatellite annotation thresholds relative to other microsatellite annotation tools. We further optimized TRF’s parameter thresholds, which we first briefly describe. TRF operates by k-tuple matching of short strings (termed ‘probes’) that slide across the reference sequence, and it evaluates matches based on probe occurrence history. TRF uses three sets of parameters to identify microsatellites: alignment parameters, conservation parameters, and a selection parameter. The alignment parameters define thresholds for how well a candidate microsatellite of length *n* must match a perfect repetition of the microsatellite motif of length *m*. Specifically, a numerical weight is assigned to each matching base (matching weight), and a numerical penalty is assigned to each mismatch or insertion/deletion (indel) position (mismatch penalty and indel penalty, respectively). The alignment score of the candidate microsatellite locus is then calculated by summing the matching weights and subtracting the mismatch and indel penalties for each position. If the alignment score exceeds a user-defined threshold, the candidate microsatellite locus is output by TRF. This minimum alignment score also indirectly sets the minimum length of annotated microsatellites since a perfect microsatellite of length *n* would have an alignment score of *n* x matching weight. This minimum length, together with the maximum period size parameter (see below), determines the minimum number of repeats required for loci with the maximum period size.

The conservation parameters are a second set of TRF filters that specify the minimum acceptable average percent identity between all adjacent copies of a motif in a candidate microsatellite (match probability) and the maximum acceptable average percentage of insertions and deletions between all adjacent copies of a motif in a candidate microsatellite (indel probability). These parameters help distinguish directly adjacent microsatellites with slightly different motifs as distinct microsatellites. Additionally, these parameters cause candidate microsatellites sharing the same motif but separated by bases that do not match the motif to be called as distinct microsatellites, rather than as a single microsatellite with a large percentage of indels (which may otherwise be filtered altogether due to the indel penalty and alignment score parameters). The selection parameter is the last TRF filter that specifies the maximum period, i.e., maximum length of the motif. Because we aimed to capture microsatellites with higher mutation rates, we set the maximum period size to 6.

To determine settings for all other TRF parameters, we reviewed settings used by 13 prior studies (Benson 1999; Gardner et al. 2002; Denœud and Vergnaud 2004; Boby et al. 2005; Domaniç and Preparata 2007; De Grassi and Ciccarelli 2009; Ryba et al. 2011; Willems et al. 2016; Bilinski et al. 2017; Gymrek et al. 2017; Karimzadeh et al. 2018; Marchal et al. 2018) and the settings recommended by the developer of TRF (Benson 1999). Based on this, we decided to use a matching weight of 2, since this was used universally across all surveyed studies and because we reasoned that annotating microsatellite would depend more on the difference between the matching weight and the mismatch and indel penalties than it would depend on the absolute value of the matching weight. Consequently, we set our minimum alignment score to 36, since this would set the minimum total microsatellite length to 18 base pairs (bp) and the minimum number of repeats of 3 for loci with the maximum period size of 6. We also used 80 and 10 for our match and indel probabilities, respectively, as these settings were used in nearly all surveyed studies.

The parameters that varied among surveyed studies were the mismatch and indel penalties. Lower penalty values increase detection of smaller microsatellites with more imperfect repeats. To find the best parameters for identifying as many microsatellites as possible while excluding loci with highly imperfect motif repeats, we tested TRF with 36 combinations of mismatch and indel penalty values, from least stringent (mismatch and indel penalty both = 3) to most stringent (mismatch and indel penalty both = 20). We evaluated the generated datasets for total microsatellite count, microsatellite lengths, average percent match, and average percent indels.

Analysis of these datasets resulted in three major conclusions: 1) indel penalty tends to have a stronger effect on the overall percent match (a measure of the “perfection”) of a given microsatellite than mismatch penalty (**Figs. S1B-C**); 2) higher mismatch and indel penalties yield shorter and more perfect microsatellites (**Figs. S1B-C**); and 3) very high mismatch penalty (MP) and indel penalty (IP) values (MP = 20 and IP = 20) do not yield major improvements in microsatellite quality (i.e., more perfect repeats) over strict, but not as high penalty values (e.g., MP = 7 and IP = 7), and they lead to exclusion of many microsatellites that should not be excluded prior to further analyses, as these microsatellites may still have elevated mutation rates despite having some imperfect repeats (**Figs. S1B-C**). Additionally, the least stringent mismatch and indel penalties (MP = 3 and IP = 3) yielded a large number of microsatellites that had very high numbers of indels and mismatches that would need to be excluded in any future filtering steps due to very low mutation rates, and thus we should not use these settings. Further analyses showed that frequencies of different motifs out of all annotated loci were stable across parameter values. Additionally, more lenient parameter values did not significantly change the length or motif distribution of the pool of returned microsatellites (**Fig. S1C**). Because the use of more lenient parameters provided no clear benefit in terms of the number, length, or types of microsatellites identified, we decided to use the values recommended by the TRF manual for these parameters (i.e., MP = 7 and IP = 7). Our final TRF parameters were: matching weight = 2, mismatch penalty = 7, indel penalty = 7, match probability = 80, indel probability = 10, minimum alignment score = 36, maximum period length = 6.

#### Microsatellite feature annotations

TRF annotates each microsatellite with several features: chromosome, start/end position, period size, copy number (i.e., the number of repeats of the microsatellite motif), consensus motif (i.e., the best-fitting motif of the microsatellite as identified by TRF), consensus size (i.e., the length of the consensus motif, which may or may not be equal to the period size), percent match, percent indel, alignment score (i.e., the sum of the matching weight, mismatch penalty, and indel penalty), % A/C/G/T, entropy (i.e., the balance of the base composition of the microsatellite; a microsatellite with 25% A, 25% C, 25% G, and 25% T has maximal entropy), and microsatellite sequence.

We also added many additional annotations to these data as follows. We derived the microsatellite motif from the TRF consensus motif, because for some loci, the TRF consensus motif was > 7 bp, despite the fact that the maximum period size was set to 6 (this has to do with how TRF works, as it defines consensus motif as the motif that maximizes the alignment score of the called microsatellite, rather than simply using the motif that the program used to call the microsatellite in the first place). We excluded microsatellites with a consensus size > 7, and for consensus motifs with a size of 7, we compared it to all possible 6 bp motifs using the adist function in R and changed it to the most similar motif. We did this because our definition of a microsatellite included a maximum motif size of 6 bp. After defining the microsatellite motif, we added an annotation for motif family, which is the motif after collapsing all circular permutations (e.g., ACG, CGA, GAC) and reverse complements of the microsatellite motif and its permutations (e.g. the reverse complement of ACG would be CGT, and then its circular permutations would be CGT, GTC, TCG) into a single representative motif family (e.g., the “ACG” motif family includes microsatellites with any of the following motifs: ACG, CGA, GAC, CGT, GTC, and TCG). We also excluded microsatellites that have another microsatellite overlapping > 90% of its span. Next, we added annotations for uninterrupted length (i.e., the length, in bp, of the longest span of perfect repeats of the microsatellite base motif without any indels or mismatches) and uninterrupted copy number (i.e., the maximum number of consecutive perfect repeats of the base motif without any indels or mismatches), as these parameters have been shown to positively correlate with the microsatellite mutation rate (Hutter et al. 2006; Gupta et al. 2007). We also annotated the 150 bp flanking sequences of the microsatellites, the overlap between each microsatellite and its neighboring microsatellite(s), the distance to the next nearest microsatellite in both 5’ and 3’ directions, and the percent GC content and mappability (using 24, 36, 50, and 100 k-mers) (Madsen et al. 2008) for each microsatellite and its 5’ and 3’ flanking regions. We further annotated the coverage depth (normalized to genome-wide average coverage) for each microsatellite and its flanking regions in a prior single-cell sequencing dataset amplified by Primary Template-directed Amplification (PTA, Bioskryb) (Gonzalez-Pena et al. 2021) in anticipation of potential future design of panels for profiling single cell genomes that have undergone initial non-uniform single-cell genome amplification. We also added annotations for replication timing based on 4D nucleome data in GM12878 lymphoblastoid cells as well as Repli-chip data of neural progenitor cells (Mizuta et al. 2004; Mousavi et al. 2019). Finally, we annotated estimated mutation rates calculated by a linear regression model we created with the ‘lm’ function in R (with default settings) trained on mutation rate data of autosomal intergenic microsatellite loci from a prior study (Gymrek et al. 2017); the model specifically utilized the ‘uninterrupted length’ and ‘motif size’ annotations to predict mutation rate, because these best explained the variance in the test dataset (Gymrek et al. 2017).

#### Building STRATIFY

We built a user-friendly, web-based app that allows users to filter microsatellites based any combination of the above annotations (**Fig. S1D**). It also dynamically updates several plots for data visualization and quality control (e.g., histogram of common microsatellite motifs by length and histogram of estimated mutation rates). Once the desired filters are selected, the microsatellite panel can be exported along with any desired annotations. The app utilizes the Shiny platform (Chang et al. 2023) along with the following packages for R statistical software version 4.2.3 (Team 2010): memoise, shiny, shinyjs, shinyWidgets, shinybusy, shinyBS, shinyalert, dplyr, readr, data.table, ggplot2, GenomicRanges. See the “Code Availability” section for a link to the full open-source code and instructions for setting up STRATIFY.

### Probe panel design

#### Pilot panel

We used STRATIFY to design the pilot hybridization capture panel testing 3 different probe design strategies: strategy 1 with only one flank of the microsatellite targeted by a 120 bp probe, strategy 2 with both flanks of the microsatellite targeted — each by a separate 120 bp probe, and strategy 3 with both flanks of the microsatellite targeted by a single 120 bp probe (half the probe complementary to the 5’ flank and the other half complementary to the 3’ flank). Using STRATIFY’s adjustable filters, we iterated systematically through combinations of parameters to find filtering thresholds for each strategy that removed unsuitable loci and left at least 100,000 loci for possible targeting (**Fig. S3B**). Once we determined the optimal filtering thresholds for each strategy separately, we combined these filters to accommodate all three design strategies.

The strictness of the filtering for the combination of strategies was largely determined by strategy 3, because both flanks needed to pass the thresholds and because we needed to use a stricter flank mappability filter since only 60 bp from each flank would be in the probe design and shorter sequences typically have lower mappability scores (**Fig. S3A**). The full set of requirements for loci in the pilot panel was: a) total microsatellite length ≤ 100 bp; b) period size between 2 to 4 bp; c) percent match ≥ 90%; d) percent indel ≤ 10%; e) uninterrupted copy number ≥ 4; f) distance to the nearest microsatellite in both directions ≥ 100 bp; g) GC content in both the microsatellite and its flanks ≤ 70%; h) 50-kmer mappability score in both 90 bp flanks ≥ 0.95 (note: the 50-kmer mappability was used to ensure uniqueness of the 60 bp flanking regions targeted by strategy 3); i) single-cell PTA normalized coverage (see above) between 0.4 and 2 in both flanks to target more uniformly amplified regions that may be useful in future single-cell applications; j) neither 90 bp flank overlaps either the 1000 Genomes Project pilot accessibility mask (regions that are challenging to resolve by short-read sequencing) (Auton et al. 2015) or the ENCODE Data Analysis Center (DAC) Blacklist Regions (regions that are prone to artifacts and erroneous signal in short-read sequencing) (Amemiya et al. 2019); h) neither 90 bp flank or the microsatellite overlap annotated segmental duplications (Bailey et al. 2002) (**Fig. S3A**). 102,456 loci passed these filters and were submitted to Twist Bioscience for probe design and synthesis.

Twist Bioscience applied proprietary quality filters to the full set of loci and assembled a list of approximately 83,000 loci that passed Twist’s filters in both flanks. From these loci, we randomly assigned 3,333 loci to each design strategy, for a total of 9,999 loci in the panel (**Table S1**). The probes were then designed as described for each strategy and delivered as a single, combined panel with each probe present at an equal concentration.

#### Large-scale panel

We used STRATIFY to design the large-scale hybridization capture panel, which targeted microsatellites with either strategy 1 (only one flank passes filters) or strategy 2 (when both flanks pass filters). The full set of requirements for loci in the large-scale panel was: a) period size between 2 to 4 bp; b) percent match ≥ 90%; c) percent indel ≤ 10%; c) uninterrupted copy number ≥ 4; d) GC content in both the microsatellite and its flanks ≤ 70%; e) distance to the nearest microsatellite ≥ 100 bp for at least one direction; f) 100-kmer mappability score in at least one of the two 90 bp flanks ≥ 0.95; g) single-cell PTA normalized coverage between 0.4 and 2 in both flanks; h) maximum of one 90 bp flank overlaps either the 1000 Genomes Project pilot accessibility mask (Auton et al. 2015) or the ENCODE Data Analysis Center Blacklist Regions (Amemiya et al. 2019); i) at least one 90 bp flank and the microsatellite itself do not overlap annotated segmental duplications (Bailey et al. 2002); j) at least one 90 bp flanks and the microsatellite itself do not overlap pseudoautosomal regions (**Fig. 3A**). Importantly, we required for each microsatellite locus that the mappability filters, region filters, and distance to the nearest microsatellite filter all pass in the same flank (for at least one flank). 173,996 loci passed these filters. Note, for the large-scale panel, we removed the filter for maximum microsatellite length, because the number of repeat units in any sample may vary from the reference genome, and because only 3.2% of microsatellites are longer than the 100 bp length limit used for the pilot panel.

We separately selected poly-A loci on chromosome X using the same filters as for the above large scale panel of 2 to 4 bp microsatellites, but with the following filters removed: a) percent match and percent indel, which are likely to be lower given the mutability of poly-A loci; b) uninterrupted copy number, because poly-A loci are very mutable and therefore unlikely to match uninterrupted copy number in the reference genome; c) GC content of the microsatellite, because poly-A loci have a GC content ∼ 0%; d) GC content of the flanks, in order to retain more loci. 9,163 poly-A loci passed these filters.

We submitted the 173,996 loci with 2 to 4 bp motifs and the 9,163 poly-A loci to Twist Biosciences, who applied final proprietary filters for probe design and synthesis. Twist was able to design probes for 147,656 loci with 2 to 4 bp motifs and for 6,532 poly-A loci by targeting at least one flank (**Table S4**). Of these, 74,635 loci with 2 to 4 bp motifs and 1,205 poly-A loci were targeted by probes in both flanks (strategy 2) and the rest were targeted by a probe in only one flank (strategy 1).

### Sample sources

Genomic DNA from lymphoblastoid cell lines was obtained from the Coriell Institute. Genomic DNA for Family 1100 and Family 2100 was extracted with the MagAttract HMW DNA Kit (Qiagen) from whole blood samples of individuals enrolled in a human subjects research protocol approved by the New York University Grossman School of Medicine Institutional Review Board.

### Library preparation

Genomic DNA was fragmented on an LE220 instrument (Covaris) using a 96 microTUBE Plate (Covaris) in a volume of 55 μL with a buffer of 10 mM Tris pH 8. Instrument settings were: 450 W peak incident factor, 20% duty factor, 1000 cycles per burst, and a total treatment time of 200 seconds. Fragmented DNA was profiled on a High Sensitivity D1000 ScreenTape TapeStation Assay (Agilent). Fragmented DNA ranged in size from 50 bp to 500 bp.

Next, we performed a nick ligation reaction to seal nicks caused by fragmentation, which may otherwise allow library preparation polymerases to initiate synthesis of new DNA strands with stutter artifacts. Nick ligation was performed in a 32 μL reaction containing 500 ng of fragmented genomic DNA, 3.73 μL NEBNext Ultra II End Prep Reaction Buffer (NEB), 26 μM NAD+, and 0.5 U/ μL E. coli DNA Ligase (NEB). The reaction was incubated at 16 °C for 30 minutes. For samples with lower concentrations (i.e., blood samples), the nick ligation reaction was scaled up to 65 µL to accommodate a larger volume of fragmented DNA.

End repair and adaptor ligation steps of library preparation were performed with the NEBNext Ultra II DNA Library Prep Kit for Illumina (NEB) at either the standard total reaction volume of the manufacturer’s protocol, or in a reaction volume scaled to 117% of the manufacturer’s protocol for samples for which the nick-ligated DNA had low concentration. The standard volume (60 μL) end repair reaction was performed by combining nick-ligated DNA, 3.27 μL NEBNext Ultra II End Prep Reaction Buffer (NEB), 3 μL NEBNext Ultra II End Prep Enzyme Mix (NEB), and nuclease-free water (NFW). The reaction was incubated at 20 °C for 30 minutes and then at 65 °C for 30 minutes. Subsequently, the standard volume (93.5 μL) adaptor ligation was performed by combining 2.5 μL of 15 μM xGen UDI-UMI adaptors (IDT), 30 µL NEBNext Ultra II Ligation Master Mix, and 1 µL of NEBNext Ligation Enhancer, then incubating at 20 °C for 15 minutes. The xGen UDI-UMI adapters contain both unique dual indexes (UDIs) for identifying the sample and unique molecular identifiers (UMIs) for identifying copies of the same molecule.

After ligation, a double-sided size selection was performed with SPRI beads (solid-phase reversible immobilization; made by washing 1 mL Sera-Mag SpeedBead Carboxylate-Modified [E3] Magnetic Particles (Cytiva) and resuspending the beads in 50 mL of 18% PEG-8000, 1.75 M NaCl, 10 mM Tris pH 8, 1 mM EDTA, 0.044% Tween-20) (Abascal et al. 2021). For the first bead addition, a ratio of 0.4X bead volume to sample volume was used, and the supernatant was retained. For the second bead addition, a bead volume of 0.25X the original volume (before the first bead addition) was used, and the sample was eluted in 22 μL of 10 mM Tris pH 8.0.

### Pre-hybridization library amplification

#### Pre-hybridization low-temperature amplification

Pre-hybridization low-temperature amplification was performed in a 50 μL reaction volume containing 20 μL ligated library, 25 mM Tris pH 7.5, 40 mM NaCl, 6 mM MgCl2, 1 mM DTT, 2 μM P5 primer (see sequence in “Library amplification primers” section), and 1 mM dNTP mix. Before the reaction, Sequenase 2.0 DNA Polymerase (Thermo Fisher) was diluted in a buffer containing 25 mM Tris pH 7.5, 40 mM NaCl, and 1 mM DTT to a final polymerase concentration of 0.67 U/μL.

Libraries underwent initial denaturation on a thermal cycler at 98 °C for 30 s, then left on the thermal cycler for the denaturation step of the first cycle for another 20 s. Samples were then immediately transferred to a water bath at 34 °C for annealing for 20 s. Excess water was quickly blotted with a Kimwipe and the samples were briefly spun down in a desktop centrifuge to remove any remaining water. Samples were then rapidly transferred to a thermal cycler set to 34 °C for extension.

Immediately after transfer to the extension thermal cycler, 0.75 μL of diluted polymerase (0.5 U) was added to the reaction, the sample was pipette mixed, and the tube was recapped with new caps since the seal created by the caps weakens after heat exposure, which can allow water to enter the reaction during annealing in the water bath. Extension continued for 90 s after polymerase was added. In the third cycle, after the extension incubation was completed, 1 μL of 100 μM P7 primer (see sequence in “Library amplification primers” section) was added to the reaction and pipette mixed to initiate the exponential phase of the reaction. After extension, the samples were transferred back to the 98 °C thermal cycler to complete the cycle (**Figs. 2A-B**). The above cycling process was repeated for 3 linear cycles and 2 exponential cycles, for a total of 5 cycles. After the last cycle’s extension step, samples were left on the 34 °C thermal cycler for a final 5 minute extension. Libraries were then purified with a 1X SPRI bead cleanup with two 80% ethanol washes and eluted in 22 μL of 10 mM Tris pH 8.0. Final concentrations were measured with the Qubit 1X dsDNA HS Assay Kit (Thermo Fisher), and the libraries were profiled with a TapeStation HS D1000 ScreenTape Assay (Agilent).

#### Pre-hybridization standard-temperature amplification

Pre-hybridization standard-temperature library amplification was performed in a 50 μL PCR reaction containing 20 μL ligated library, 1X NEBNext Ultra II Q5 Master Mix (NEB), and 2 μM P5 primer. Thermal cycling was conducted as follows: initial denaturation at 98 °C for 30 s, denaturation at 98 °C for 10 s, annealing and extension at 65 °C for 75 s, final extension at 65°C for 5 minutes (**Fig. 2B**). P7 primer was added after the extension step of the third cycle as in the low-temperature amplification to start the exponential cycles. The cycling process was repeated for 3 linear cycles and 2 exponential cycles, as was performed for the low-temperature pre-hybridization amplification. The samples were purified, quantified, and profiled as described for the low-temperature amplification.

#### Library amplification primers

Library amplification primers were ordered as HPLC-purified oligonucleotides from Integrated DNA Technologies (IDT).

P5-PTx2: AATGATACGGCGACCACCGA*G*A

P7-PTx2: CAAGCAGAAGACGGCATACG*A*G

* phosphorothioate bonds

### Hybridization capture

Hybridization capture was performed with the Standard Hybridization Reagent Kit v2 (Twist Bioscience) and biotinylated 120 bp probes delivered at a concentration of 0.35 fmol/probe (Twist Bioscience) per the manufacturer’s protocol. In pilot panel hybridizations, 375 ng of each library was input into a 4-plex hybridization. In large-scale panel hybridizations, 350 ng (Coriell 1, Coriell 2, Coriell 3 trios) or 250 ng (Family 1100 and Family 2100 trios) of each library were input into the hybridizations. Each Coriell trio was hybridized in a separate 3-plex hybridization, and Family 1100 and Family 2100 trios were multiplexed together in a single hybridization.

### Post-hybridization library amplification

#### Post-hybridization low-temperature amplification

Post-hybridization low-temperature amplification was performed in a 50 μL reaction volume containing 22.5 μL of the streptavidin bead slurry from hybridization, 1X NEBuffer 2 (NEB), 0.5 μM P5 primer, and 1 mM dNTP mix. Klenow Fragment (3’→5’ exo-) (NEB) was diluted in 1X NEBuffer 2 to a final concentration of 2 U/μL. Thermal cycling and polymerase addition was conducted as in the pre-hybridization low-temperature amplification. Note that post-hybridization amplification utilized diluted Klenow Fragment polymerase rather than the Sequenase 2.0 polymerase used in the pre-hybridization amplification. After the extension step of the third cycle, 1 μL of 25 μM P7 primer was added to the reaction and pipette mixed to start the exponential phase of the reaction (**Figs. 2A-B**). The cycling process was repeated for 3 linear cycles and 8 exponential cycles for the pilot panel, and 3 linear cycles and 6 exponential cycles for the large-scale panel. The samples were purified with a 1X DNA purification bead (Twist Bioscience) cleanup with two 80% ethanol washes and eluted in 25 μL of 10 mM Tris pH 8.0. Since the 3’→5’ exo-Klenow Fragment leaves a 3’ non-templated nucleotide at the ends of molecules (Clark et al. 1987), we removed these in a 35 μL reaction containing the purified library, 1X NEBuffer 2, 0.033 mM dNTP mix, and 1 U of DNA Polymerase I, Large (Klenow) fragment (NEB), incubated at 25 °C for 15 minutes. The reaction was stopped by adding EDTA to a final concentration of 10 mM and purified with a 1X DNA purification bead (Twist) cleanup with two 80% ethanol washes and eluted in 32 μL of 10 mM Tris pH 8.0.

#### Post-hybridization standard-temperature amplification

Post-hybridization standard-temperature library amplification was performed in a 50 μL PCR reaction containing 22.5 μL of the streptavidin binding bead slurry from hybridization, 0.5 μM P5 primer, and 1X Equinox Library Amp Master Mix (Twist Bioscience). The thermal cycling protocol was as follows: initial denaturation at 98 °C for 45 s, denaturation at 98 °C for 15 s, annealing at 60 °C for 30 s, extension at 72 °C for 30 s, final extension at 72 °C for 60 s (**Fig. 2B**). P7 primer was added after the extension step of the third cycle as in the low-temperature amplification. The cycling process was repeated for 3 linear cycles and 8 exponential cycles for the pilot panel, as was performed for the low-temperature post-hybridization amplification. The libraries were purified, quantified, and profiled as in the low-temperature post-hybridization amplification.

### Sequencing

Sequencing was performed on NovaSeq 6000 instruments (lllumina) at the University of California Irvine Genomics High-Throughput Facility and the University of California Berkeley QB3 Genomics. See **Tables S2** and **S5** for sequencing metrics for all samples in the study.

### Computational pipeline for microsatellite genotyping

#### Molecular duplication analysis

Molecular duplication rates were calculated for samples from trios for which unique molecular identifiers (UMI) were sequenced (Family 1100 and Family 2100). The computational pipeline (schematic in **Fig. S6B**) converted FASTQ sequencing files to unmapped BAMs with the fgbio v2.0.2 (Fennell and Homer) FastqToBam tool with the option --extract-umis-from-read-names to extract UMI sequences. The unmapped BAM file was then reverted to FASTQ format with samtools v0.11.9 fastq (Danecek et al. 2021), then aligned to the reference genome with BWA-MEM v0.7.17 (Li 2013) using options “-p -K 150000000 -Y”. The mapped BAM and the unmapped BAM were input into fgbio ZipperBams (Fennell and Homer) to add the UMI metadata to the mapped BAM, and the mapped BAM was subsequently sorted by the query name with the samtools v1.14 tool sort (Danecek et al. 2021) using the option “-n”. After sorting, optical duplicates were removed (since these should not be included in the UMI metrics as they are not true molecular duplicates) with the Picard v2.27.4 MarkDuplicates tool (Institute 2019) with options “REMOVE_DUPLICATES=false OPTICAL_DUPLICATE_PIXEL_DISTANCE=2500 ASSUME_SORT_ORDER=queryname CLEAR_DT=false REMOVE_SEQUENCING_DUPLICATES=true READ_NAME_REGEX=[a-zA-Z0-9]+:[0-9]+:[a-zA-Z0-9]+:[0-9]:\([0-9]+\):\([0-9]+\):\([0-9]+\).*”. The BAM file was then unmarked with the Genome Analysis Toolkit (GATK) v4.2.4.1 (McKenna et al. 2010; DePristo et al. 2011; Van der Auwera et al. 2013) UnmarkDuplicates tool. Finally, the number of UMIs observed for each molecular duplicate family size (where family size is the number of molecules observed with the same UMI sequence) was calculated using fgbio GroupReadsByUmi (Fennell and Homer) with options “--strategy Adjacency --edits 1 --family-size-histogram”.

#### STREAM ensemble microsatellite genotyping pipeline

STREAM, our ensemble microsatellite genotyping pipeline is run via the Nextflow v21.10.6 scientific workflow tool (Di Tommaso et al. 2017) (see schematic in **Figs. S6C-D**). Reads were aligned to the human reference genome (hg38) with BWA-MEM v0.7.17 (Li 2013). Optical duplicates were removed with Picard MarkDuplicates (Institute 2019) with options “REMOVE_DUPLICATES=false OPTICAL_DUPLICATE_PIXEL_DISTANCE=2500 ASSUME_SORT_ORDER=queryname CLEAR_DT=false REMOVE_SEQUENCING_DUPLICATES=true READ_NAME_REGEX=[a-zA-Z0-9]+:[0-9]+:[a-zA-Z0-9]+:[0-9]:\([0-9]+\):\([0-9]+\):\([0-9]+\).*” (note: the read name regular expression was included to handle files with UMI sequences). The fraction of optical duplication was calculated with the equation (READ_PAIR_OPTICAL_DUPLICATES x 2) / (UNPAIRED_READS_EXAMINED + READ_PAIRS_EXAMINED x 2), based on MarkDuplicates documentation. The resulting BAM file was unmarked using GATK UnmarkDuplicates (McKenna et al. 2010; DePristo et al. 2011; Van der Auwera et al. 2013) and sorted with Picard SortSam with parameter “SORT_ORDER=coordinate”, followed by conversion to CRAM format with samtools v1.14 view (Danecek et al. 2021) with options “-F 2304 -C” (filters out non-primary and supplementary alignments). Hybridization capture quality metrics — including percent of on-target reads, percent of zero-coverage loci, AT-dropout, and fold 80 base penalty — were calculated using Picard CollectHsMetrics (Institute 2019).

The aligned reads were then input into 3 different microsatellite callers: HipSTR v0.7 (Willems et al. 2017), GangSTR v2.5.0 (Mousavi et al. 2019), and ExpansionHunter v5.0.0 (Dolzhenko et al. 2019). HipSTR was run on all samples jointly with parameters “--min-reads 20 --max-str-len 150 --no-rmdup --output-filters”, and the output VCF was split by sample using BCFtools (Danecek et al. 2021). For the temperature-comparison assay, HipSTR was also run with the option “--haploid-chrs chrX,chrY” because all of the replicates being profiled were male. GangSTR and Expansion Hunter do not support joint genotyping and were run on each sample individually. GangSTR was run with parameters “--min-sample reads 20 --nonuniform --frrweight 0 --spanweight 0 --flankweight 0”, and ExpansionHunter was run with default parameters. For both GangSTR and ExpansionHunter, each sample’s sex was specified. Since ExpansionHunter does not output total read depth for loci, which is required for downstream analysis of ExpansionHunter data, the bedtools coverage tool (Quinlan and Hall 2010) was used with parameters “-sorted -f 1.0” to calculate the number of mapped reads that completely span the coordinates of the microsatellite loci (spanning reads). Relevant fields were then extracted from each caller’s VCF output and converted to a tab-delimited format with BCFtools (Danecek et al. 2021).

Genotype information from each caller was further processed with an R (Team 2010) script (requiring the packages dplyr (Wickham et al. 2023a) and tidyr (Wickham et al. 2023b)), as follows:

1. The capture panel information, sample information, and data from HipSTR, GangSTR, ExpansionHunter, and bedtools coverage analyses are loaded and joined into a single table.
2. ExpansionHunter calls for which the bedtools spanning read coverage is 0 are excluded from further analysis.
3. HipSTR diploid calls for chromosome X loci in male samples are converted to haploid calls by keeping only the allele with the most supporting reads per the MALLREADS field.
4. For each caller, we calculate the difference in repeat units of the called genotype from the reference genome.
5. For HipSTR only, we calculate the fraction of total reads (DP) with stutter (DSTUTTER) and the fraction of total reads with flank indels (DFLANKINDEL).
6. For HipSTR, we sum the reads reported in the MALLREADS field to obtain the total number of reads used to call the genotype at each locus.
7. For GangSTR, we sum the reads reported in the ENCLREADS field to obtain the total number of spanning reads reported by GangSTR.
8. The variant allele fraction (VAF) of each allele is calculated by dividing the number of reads supporting that allele by the total number of reads at the locus, as follows:

a. For HipSTR, read classes are not reported separately, so we use the number of reads in the MALLREADS field that support the called allele as the numerator and the total number of reads reported in MALLREADS as the denominator.
b. For GangSTR, we use the number of reads in the ENCLREADS field that support the called allele as the numerator and the total number of reads reported in ENCLREADS as the denominator. We use the ENCLREADS because this reports only the spanning reads.
c. For ExpansionHunter, we use the number of reads in the ADSP field that support the called allele as the numerator. We use ADSP because this field reports only the spanning reads identified by ExpansionHunter. ExpansionHunter does not report a total read depth, so we use the number of spanning reads counted by bedtools coverage as the denominator (see above), which results in some VAF values greater than 1 because the bedtools coverage read count is not exactly equivalent to the reads counted by ExpansionHunter.
9. Data from all samples are joined into a single table.

At this stage, each locus has 3 genotype calls, one from each caller. For each locus, we need to choose one caller whose genotypes will be used as the final genotypes for all samples. We want to use only one caller for all samples because the reported genotype can differ between callers, precluding concordance analysis between samples if genotypes from different callers are used. The first step in choosing a caller is to filter the reported genotypes based on a variety of quality parameters (**Figs. S6D** and **S9**). The pipeline uses a YAML configuration file to store the values for each filtering threshold and an order of preference for which caller’s genotypes to use. First, sample-level filters are applied to each caller’s genotype calls for each sample. The sample-level filters common to all callers are: 1) minimum total reads (MALLREADS field for HipSTR, ENCLREADS for GangSTR, and bedtools spanning read count for ExpansionHunter); 2) minimum number of reads supporting each allele (same fields as total reads except for ExpansionHunter, which uses the ADSP field); 3) minimum VAF. ExpansionHunter has both a minimum and maximum VAF threshold due to the previously explained issue with the VAF denominator. HipSTR also has additional sample-level filters: 1) minimum quality score; 2) maximum fraction of reads containing stutter; 3) maximum fraction of reads containing flank indels; 4) minimum likelihood difference between the reported genotype and the next best genotype (GLDIFF). GangSTR also has an additional sample-level filter of minimum quality score. ExpansionHunter also has an additional sample-level filter of minimum average locus coverage. For many filters, both alleles are required to pass the filters, so the pipeline accounts for the sex of the sample when applying filters to sex chromosomes.

Next, the pipeline applies locus-level filters to check for systemic genotyping issues across multiple samples for each locus. The locus-level filters used for this step are: 1) minimum mean quality score across samples (HipSTR and GangSTR only); 2) minimum mean total read depth across samples (using the same fields as in the sample-level filters); 3) minimum fraction of samples passing the sample-level filters.

After the sample-level and locus-level filters have been applied, the final genotype call is chosen from one of the three callers. The caller used for each locus is chosen based on the caller order of preference specified in the YAML configuration file and whether the locus has passed the locus-level filters for that caller. If the locus does not pass locus-level filters for any caller, no call is reported. Then, at each locus, the pipeline checks if the sample passed the sample-level filters. If the sample passed filtering, the genotype from the caller chosen for that locus is entered into the final genotype column; if the sample did not pass filtering at that locus; no genotype is entered.

For this analysis, we chose the combination of filtering parameters and optimized their thresholds (see details of all parameters and thresholds below and **Figs. S9** and **S10**) by testing a range of parameter combinations and thresholds and calculating the Mendelian discordance rate of the resulting genotypes, i.e., the fraction of loci where the genotypes of the parents and offspring do not match patterns of Mendelian inheritance. If the genotype of the offspring does match Mendelian inheritance patterns, that locus is considered concordant. A true discordant locus indicates the presence of a de novo mutation in the offspring, so the rate of discordance should match the microsatellite de novo mutation rate, which prior studies have estimated to be ∼ 1 x 10^-4^ mutations per locus per generation (Weber and Wong 1993; Huang et al. 2002; Sun et al. 2012; Kristmundsdottir et al. 2023). In addition to optimizing our filters to achieve this discordance rate, we also optimized them to maximize the number of loci that were genotyped (i.e., that passed filters). Optimization was performed using Family 1100 and confirmed on the other trios. Mendelian discordance calculations were only performed on loci that were genotyped in all three members of the family trio (“fully genotyped loci”). For trios with a male child, loci on chromosome X were considered fully genotyped if only the mother and son were genotyped since the father did not pass on an X chromosome.

We implemented different filtering parameters and thresholds for 2 to 4 bp motif loci and poly-A loci due to the latter’s higher stutter levels. For 2 to 4 bp motif loci, we used a caller preference order of 1) HipSTR, 2) GangSTR, and 3) ExpansionHunter. For HipSTR, we used the following filter parameters: a) VAF ≥ 0.2; b) fraction of stutter reads ≤ 0.13; c) fraction of reads with flank indels ≤ 0.13; d) genotype likelihood difference ≥ 25; e) mean quality score ≥ 0.95. For GangSTR, we used the following filter parameters: a) number of reads supporting the allele ≥ 80 for both alleles; b) mean quality score ≥ 1.0. For ExpansionHunter, we used the following filter parameters: a) 0.5 ≤ VAF ≤ 2.5; b) number of reads supporting the allele ≥ 60 for both alleles.

We separately optimized filtering parameters and thresholds for poly-A loci due to their overall lower quality genotype calls and higher stutter levels. We used slightly different filters depending on whether the trio had a male or a female child because we observed that trios with female children tended to have higher discordance rates, likely because genotyping of heterozygous chromosome X loci is more challenging than genotyping hemizygous chromosome X loci. For all trios regardless of the sex of the child, we used a caller preference order of 1) GangSTR and 2) HipSTR. We did not use ExpansionHunter for poly-A loci calls because we could not find a combination of filters that achieved the expected Mendelian discordance rate. For trios with male children, we used the following filter parameters for GangSTR: a) number of reads supporting each allele ≥ 10; b) mean quality score ≥ 0.5. For HipSTR, we used the following filter parameters: a) VAF ≥ 0.2; b) fraction of reads with flank indels ≤ 0.13; c) genotype likelihood difference ≥ 1; d) mean quality score ≥ 0.85. For trios with female children, we adjusted the GangSTR filters by increasing the number of reads supporting each allele to ≥ 15 and adding a filter for VAF ≥ 0.4, and the HipSTR VAF filter was changed to ≥ 0.3.

#### Hybridization capture subsampling analysis

We performed an analysis to determine how average read depth per locus affects the results of our hybridization capture genotyping. Specifically, we compared the number of fully genotyped loci and the Mendelian discordance rate at different subsampled average read depths per locus, where average read depth per locus is the total number of aligned paired-end sequencing reads divided by the number of loci in the panel. This subsampling analysis was performed on the large-scale panel data of Family 1100 because, across all trios, its lowest coverage sample had the highest number of aligned reads. We subsampled FASTQ files for each member of the trio using seqtk sample v1.3 (Li 2012) with options “-2 -s100” to 30, 60, 120, 200, 400, 600, 800, 900, and 1013 average reads per locus. The highest subsampling level of 1013 average reads per locus corresponds to the original sample size of individual 1101, which is the sample in the trio with the lowest average reads per locus. For this subsampling level, we subsampled the other samples in the trio to the number of reads in the sample of individual 1101. For lower coverage levels, we multiplied the desired average reads per locus by the number of loci and subsampled to that number of reads. Each set of subsampled FASTQ files was processed with our STREAM pipeline along with the other four trios, whose sequencing levels had not been adjusted; we ran all samples together for consistency of HipSTR joint genotyping and genotype filtering. Finally, the Mendelian discordance rate and the number of fully genotyped loci were calculated for each subsampling level.

#### Whole genome sequencing subsampling analysis

We performed a subsampling analysis of high-depth PCR-free Illumina WGS on a family trio (HG002 = child, HG003 = father, HG004 = mother) from the NIST Genome in a Bottle (GIAB) project. We downloaded the FASTQ data from the NIST Genome in a Bottle (GIAB) (Bottle 2023). The reads were subsampled as in the hybridization capture subsampling analysis at 30X, 60X, 100X, 150X, 200X, 250X, and 309X average coverage. The highest subsampling level is the average coverage of sample HG002, which is the sample with the lowest average coverage in the trio.

Since this is WGS data, we used STRATIFY to create a genome-wide list of microsatellites for analysis, and because we no longer needed to consider capture probes in our filtering, we used the following minimal filters: a) distance to the nearest microsatellite ≥ 75 bp in both flanks; b) 100-kmer mappability score in both 90 bp flanks ≥ 0.95; c) exclusion of microsatellites for which at least one 90 bp flank overlaps either the 1000 Genomes Project pilot accessibility mask (Auton et al. 2015) or the ENCODE Data Analysis Center Blacklist Regions (Amemiya et al. 2019); d) exclusion of microsatellites for which at least one 90 bp flank and/or the microsatellite itself overlaps annotated segmental duplications (Bailey et al. 2002) or pseudoautosomal regions. We further required for each microsatellite locus that the mappability filters, region filters, and distance to the nearest microsatellite filter all pass in the same flank (for at least one flank). 563,770 loci passed these filters, 395,725 of which have 2 to 4 bp motifs and 168,045 of which have a poly-A motif.

We processed the high-depth sequencing data with Stage 1 of STREAM (**Fig. 6C**) using either the loci from the large-scale panel or the loci from the genome-wide list described above. We then re-optimized the quality filtering parameters for the 309X WGS data to obtain the expected Mendelian discordance rate for the large-scale panel. The filtering thresholds used were identical to the thresholds used for the hybridization capture large-scale panel analysis described above, except for the following filters for 2 to 4 bp motifs: a) fraction of stutter reads ≤ 0.1, since we observed slightly lower levels of stutter in the PCR-free data, and b) we turned off the filter for fraction of reads with flank indels, since we observed only a handful of loci above this threshold. The filtering thresholds used for poly-A loci were identical to those used for the hybridization capture large-scale panel analysis for trios with male children described above. Lower subsampling levels were then analyzed with STREAM using these filtering thresholds. The same filtering thresholds were used to analyze the WGS data both for the large-scale panel and the genome-wide list of loci. Finally, the Mendelian discordance rate and the number of fully genotyped loci were calculated for each subsampling level and for each set of loci.

#### Tools for data analysis and plots

We used the following data analysis tools and packages to analyze and plot our data. R packages: dplyr (Wickham et al. 2023a), tidyr (Wickham et al. 2023b), ggplot2 (Wickham 2016), gghighlight (Yutani 2022), ggbeeswarm (Clarke et al. 2023), ggforce (Pedersen 2022), wesanderson (Ram and Wickham 2018); other analysis tools: bedtools (Quinlan and Hall 2010), UCSCtools bedGraphToBigWig (Kent et al. 2010), and BioRender.com.

#### Capillary electrophoresis validation

We used capillary electrophoresis to validate discordant genotype calls in three family trios: Coriell 3, Family 1100, and Family 2100. For each locus called as discordant by STREAM, we designed a primer pair with one primer marked with fluorescein (6-FAM) at its 5’ end (see **Table S6** for primer sequences). PCR was performed in a 20 µL reaction containing 1X Colorless GoTaq Flexi Buffer (Promega), 1.5 mM MgCl2, 0.2 mM dNTP mix, 0.25 uM each primer, and 0.5 U GoTaq G2 Flexi DNA Polymerase (Promega). Thermal cycling was performed for 40 cycles with the following protocol: initial denaturation at 95 °C for 2 min, denaturation at 95 °C for 30 s, annealing at 58 °C for 30 s, extension at 72 °C for 45 s, final extension at 72 °C for 5 min. Capillary electrophoresis was performed by the University of Arizona Genetics Core. The analysis software Peak Scanner (Thermo Fisher Connect Platform) was used to determine the amplicon length for each locus and sample. Inferred repeat unit counts from the fragment analysis assay were then compared to the genotypes output by STREAM

## AVAILABILITY OF DATA AND MATERIALS

The sequencing data used in this article are available at the NCBI database of Genotypes and Phenotypes (dbGaP) [accession ID pending].

## CODE AVAILABILITY

The source code for STRATIFY is available at https://github.com/evronylab/STRATIFY. The source code for STREAM is available at https://github.com/evronylab/STREAM.

## AUTHOR CONTRIBUTIONS

C.A.L., D.S., and G.D.E. conceived the technology. C.A.L. designed the experiments with input from G.D.E. C.A.L. performed the experiments. D.S. developed STRATIFY. C.A.L. developed STREAM and analyzed the data with input from D.S., A.S., and G.D.E. C.A.L. wrote the manuscript with input from D.S. and G.D.E.

## SUPPLEMENTAL NOTE

### Cost analysis of our targeted capture method relative to whole-genome sequencing

Here, we estimate the cost of our targeted capture method, including library preparation, hybridization capture, and sequencing and compare it to the cost of PCR-free whole genome sequencing (WGS) as an alternative approach for large-scale sequencing of microsatellites. This analysis was performed by subsampling high-coverage trio data to a range of average read depths. Specifically, we utilized data from trio Family 1100 for which we have very high-read depth targeted capture data (1013X for the lowest coverage family member) and data from trio HG002/HG003/HG004 (Zook et al. 2016) with very high-read depth PCR-free WGS (309X for the lowest coverage family member). The cost estimates in the below text describe analysis of the large-scale panel of 154,188 loci in samples profiled by our method compared to analysis of a larger genome-wide list of 563,770 loci in samples profiled by WGS. The cost estimates in the below text also assume use of the Novaseq X 25B flow cell ($2 / gigabase sequenced) and exclude labor costs. In **Figs. S12A-B**, we also include cost estimates for the Novaseq 6000 S4 flow cells ($5 / gigabase sequenced) and cost estimates for analysis of the large-scale panel of loci in WGS samples.

For our targeted capture method using our large-scale panel, we estimate a cost of $127 for library preparation and hybridization per sample and $37 for sequencing per sample, assuming hybridization multiplexing of 8 samples and an average read depth per locus of 800X (i.e., 18.5 gigabases (Gb) sequencing per sample = 150 bp per read × 800 reads/locus × 154,188 loci). Therefore, the total cost is $164 per sample at this read depth. At this read depth, 84,734 loci with 2 to 4 bp motifs are fully genotyped (i.e., genotyped in all family members), yielding a cost of $1.94 per 1,000 fully genotyped locI (**Fig. S12A**).

For PCR-free WGS analysis of a genome-wide set of microsatellite loci, we estimate a cost of $22 for library preparation per sample and $600 for sequencing per sample, assuming an average read depth of 100X (i.e., 300 Gb). Therefore, the total cost is $622 per sample at this read depth. At this read depth, 271,693 loci with 2 to 4 bp motifs are fully genotyped, yielding a cost of $2.29 per 1,000 fully genotyped loci (**Fig. S12A**). These results indicate that at a similar cost per 1,000 fully genotyped loci, PCR-free WGS yields ∼ 3.2-fold more fully genotyped loci but also incurs ∼ 3.8-fold higher cost per sample relative to our targeted capture method. **Fig. S12B** shows a similar analysis for poly-A loci.

Notably, the above read depths were chosen for the comparison, because we found that at lower average read depths (< 100X for WGS and < 800X for targeted capture), the Mendelian discordance rate is lower than expected, likely because at lower read depth, non-discordant reference loci are preferentially fully genotyped by the microsatellite callers. Therefore, to reliably capture discordant loci as required by any application, these minimum read depths are needed. As a result, while WGS has potential to genotype more loci, it entails a higher minimum cost per sample. Therefore, the most cost-effective method, either targeted capture or WGS, depends on the number of loci required by the biological application.

Additionally, the total cost per sample and the cost per 1,000 fully genotyped loci of our targeted capture method will become more efficient as DNA synthesis (i.e., capture probe) costs decrease. Some applications such as lineage tracing will also benefit from very high read depth to increase the fraction of samples that are genotyped at each locus, such that our targeted capture method may be preferable because it is more cost effective relative to WGS as read depth increases (**Figs. S12A-B**). Poly-A loci in particular require high read depth to be fully genotyped (**Fig. S12B**), and a capture panel containing more poly-A loci than our current large-scale panel would achieve better cost parity with WGS at these higher read depths. Finally, some samples undergo amplification prior to profiling, such as amplified single-cell genomes, with a resulting baseline stutter that will require a higher read depth for accurate genotyping. This may be more cost effective to achieve by targeted capture than WGS, albeit while genotyping a lower total number of loci across the genome.

## SUPPLEMENTAL FIGURE AND TABLE LEGENDS

**Figure S1.**
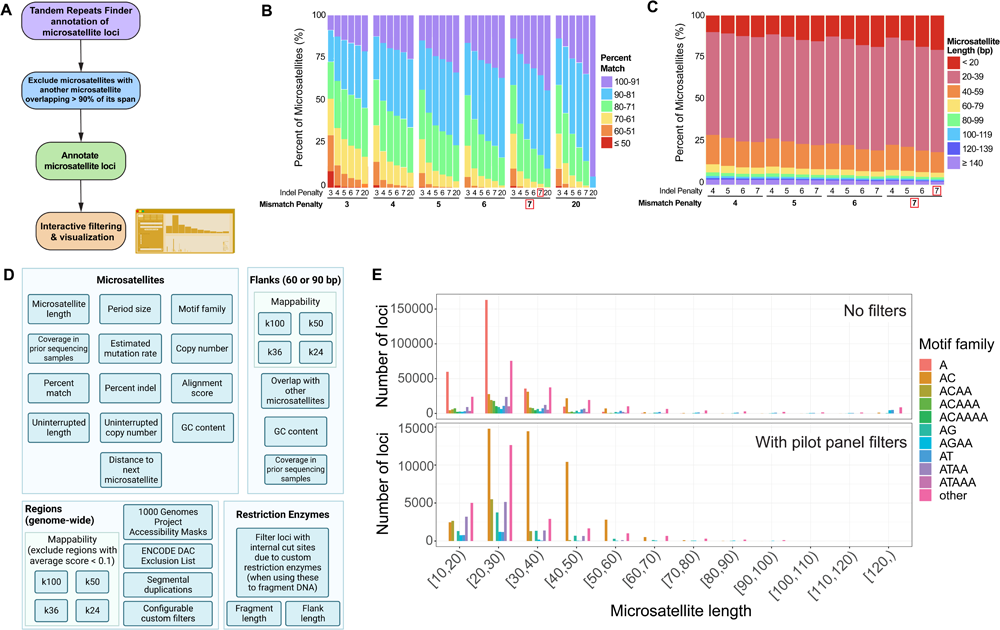
Development and design of STRATIFY. **A.** Workflow for development of STRATIFY, a web-based application for filtering and designing a panel of microsatellites for targeted hybridization capture. **B.** The effects of Tandem Repeats Finder (TRF) indel penalty and mismatch penalty parameters (i.e., numerical penalties assigned to each mismatch or indel position) on the percentage of microsatellites in different bins of TRF-calculated ‘percent match’ values (i.e., TRF-calculated percent identity between adjacent repeat units). The final settings used to generate the microsatellite dataset for STRATIFY are boxed in red. **C.** The effects of TRF indel penalty and mismatch penalty parameters on the distribution of sequence lengths of microsatellites identified by TRF. The final settings used to generate the microsatellite dataset for STRATIFY are boxed in red. **D.** All filters available in STRATIFY for refinement of a microsatellite capture panel. See **Methods** for details of each filter. **E.** Example of a plot produced by STRATIFY: histogram of the number of loci in STRATIFY before filtering (top) and after filtering for the pilot panel (bottom), binned by the length of the microsatellite and separated by motif family. The motif families shown are the ten most frequent motif families in the unfiltered dataset; all remaining loci are grouped into “other”.

**Figure S2.**
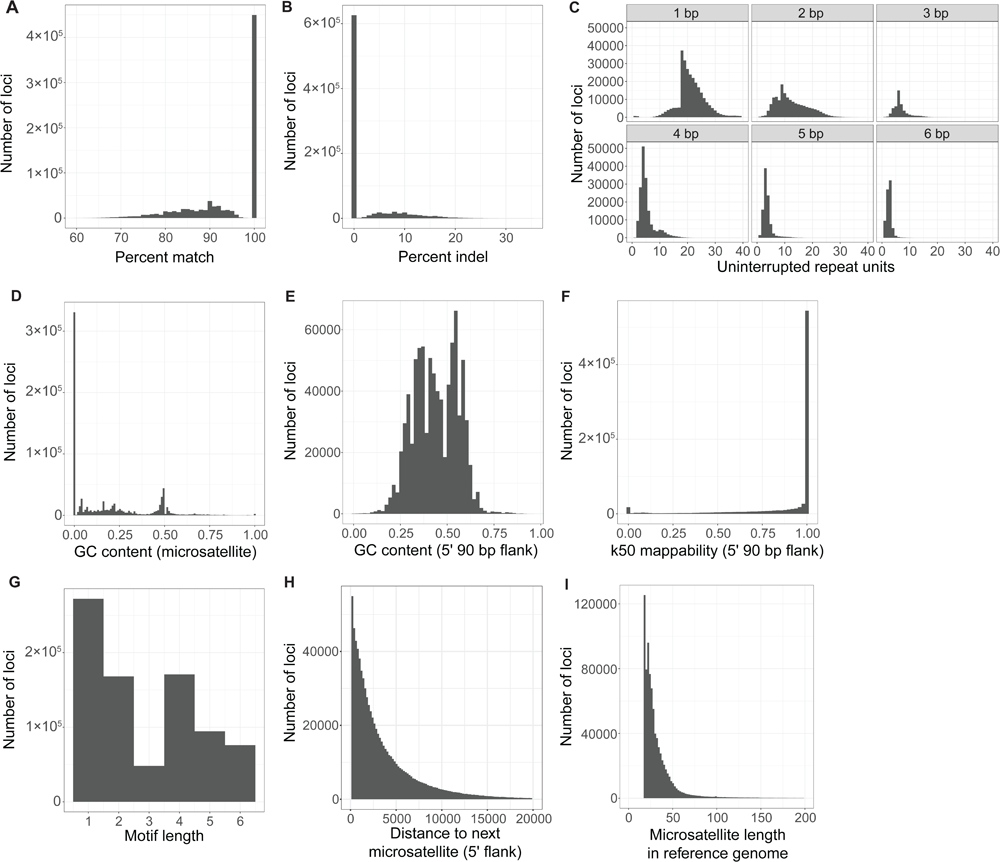
Distributions of filtering parameter values in the STRATIFY dataset. Histograms of the full STRATIFY dataset for a selection of available filtering parameters. **A.** Percent of bases in the microsatellite called as matches by TRF. **B.** Percent of bases in the microsatellite called as indels (insertion or deletion) by TRF. **C.** Longest number of consecutive repeat units uninterrupted by mismatches or indels, for different motif lengths. For shorter motifs, there are peaks in the histogram; for example, there are peaks at 18 and 9 repeat units for microsatellites with 1 bp and 2 bp motif lengths, respectively. This is because with our TRF settings of a minimum score of 36 and a match weight of 2 (**Methods**), the shortest microsatellites called by TRF are perfect repeats with a total length of 18 bp. **D.** GC content of microsatellite loci. **E.** GC content of the 5’ 90 bp flanks of microsatellite loci. The 3’ flanks have a very similar distribution. **F.** Mappability scores of the 5’ 90 bp flanks calculated using 50 bp k-mers (**Methods**). The 3’ flanks have a very similar distribution. **G.** Repeat motif lengths. **H.** Distance to the next microsatellite in the genome in the 5’ direction. Data for the 3’ direction has a very similar distribution. **I.** Microsatellite length in the reference genome (minimum 18 bp due to minimum score and match weight used in TRF, as described in **Fig. S2C**).

**Figure S3.**
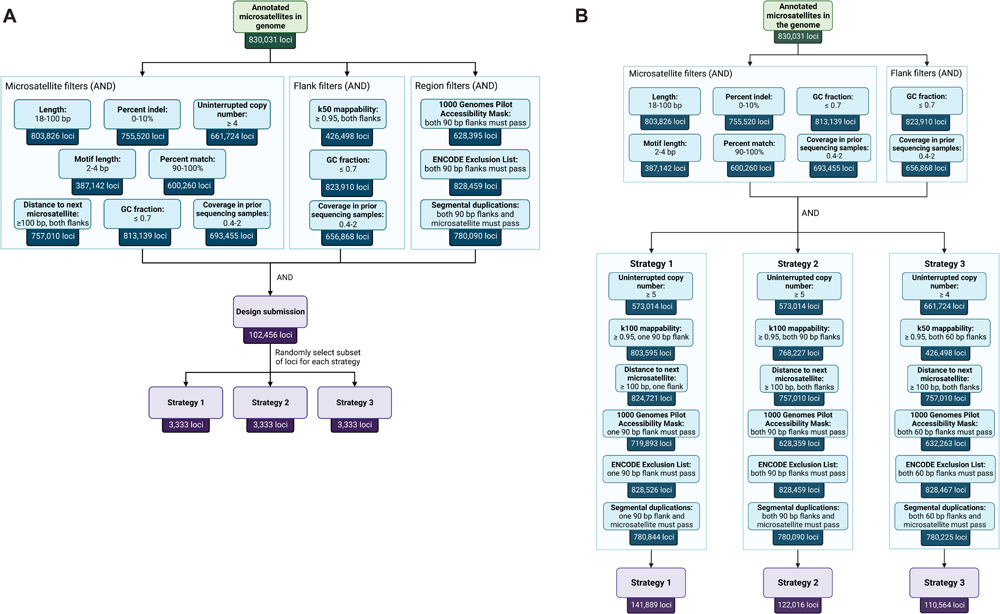
Pilot panel design. **A.** Schematic of locus selection for the pilot microsatellite capture panel that was used to test the three probe design strategies. All filters are applied with Boolean AND logic. The ‘Design submission’ step is an additional set of proprietary filters for probe synthesis and design by Twist Biosciences. See **Methods** for details of each filter. **B.** Schematic of the different filtering requirements for each probe design strategy used in the pilot panel and the final number of loci across the genome that are potentially targetable if using each strategy individually. Strategy 3 is more restrictive than Strategy 1 or 2, which would limit the size of future large-scale panels. See **Methods** for details of each filter.

**Figure S4.**
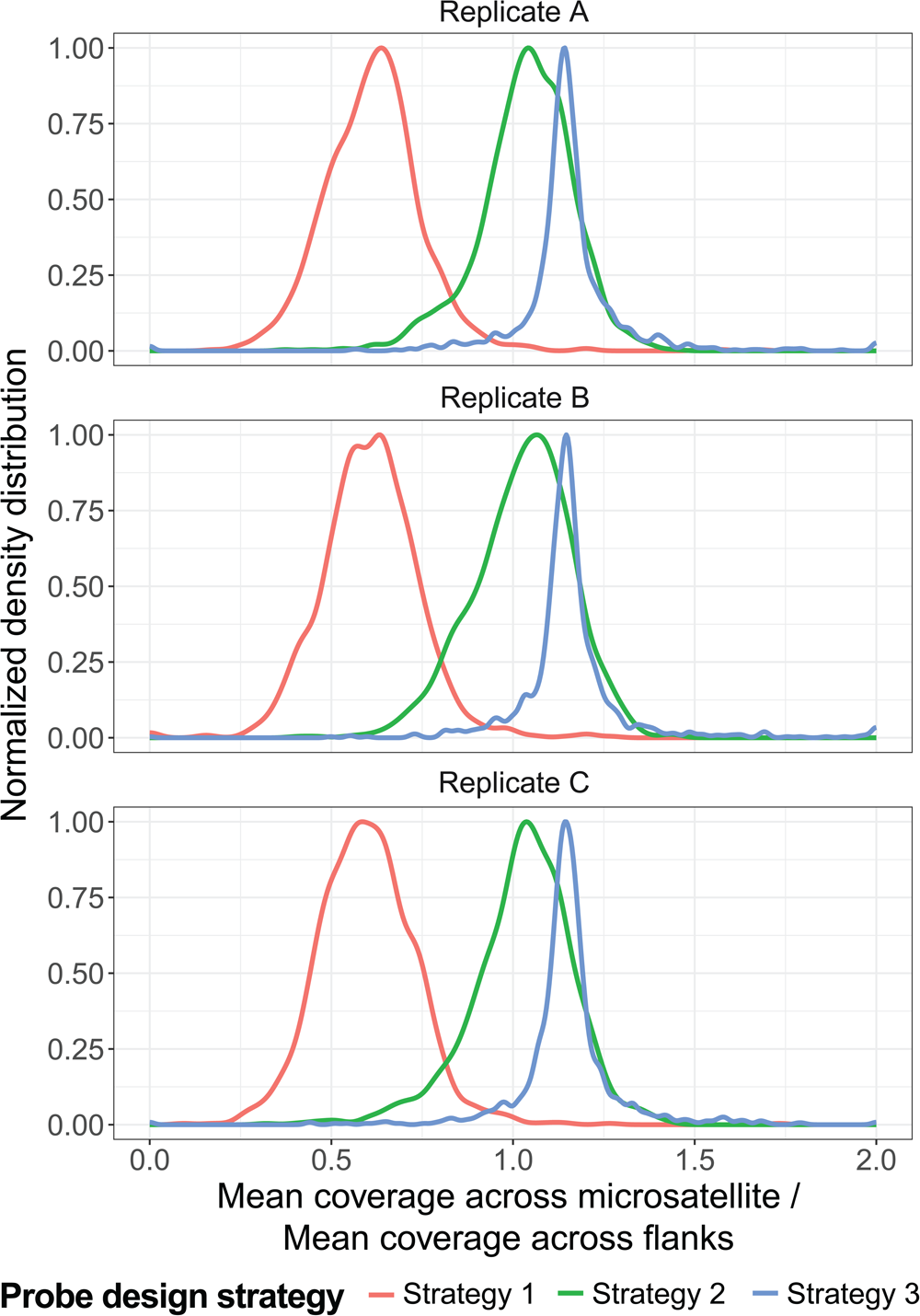
Ratio of microsatellite coverage to flank coverage for each probe design strategy. Distribution across all targeted microsatellite loci of the ratio of mean coverage (read depth) of the microsatellite to the mean coverage of its targeted flanks, for each probe design strategy. Each replicate is a technical replicate of sample NA12877 profiled with the pilot capture panel. A ratio value > 1 indicates the microsatellite had a higher mean coverage than that of the flanking regions. For Strategies 2 and 3, in which both flanks were targeted by probes, the coverage of the flanks was averaged, and this value was used to calculate the ratio. The peak of each density distribution is normalized to 1.0 for easier comparison between strategies.

**Figure S5.**
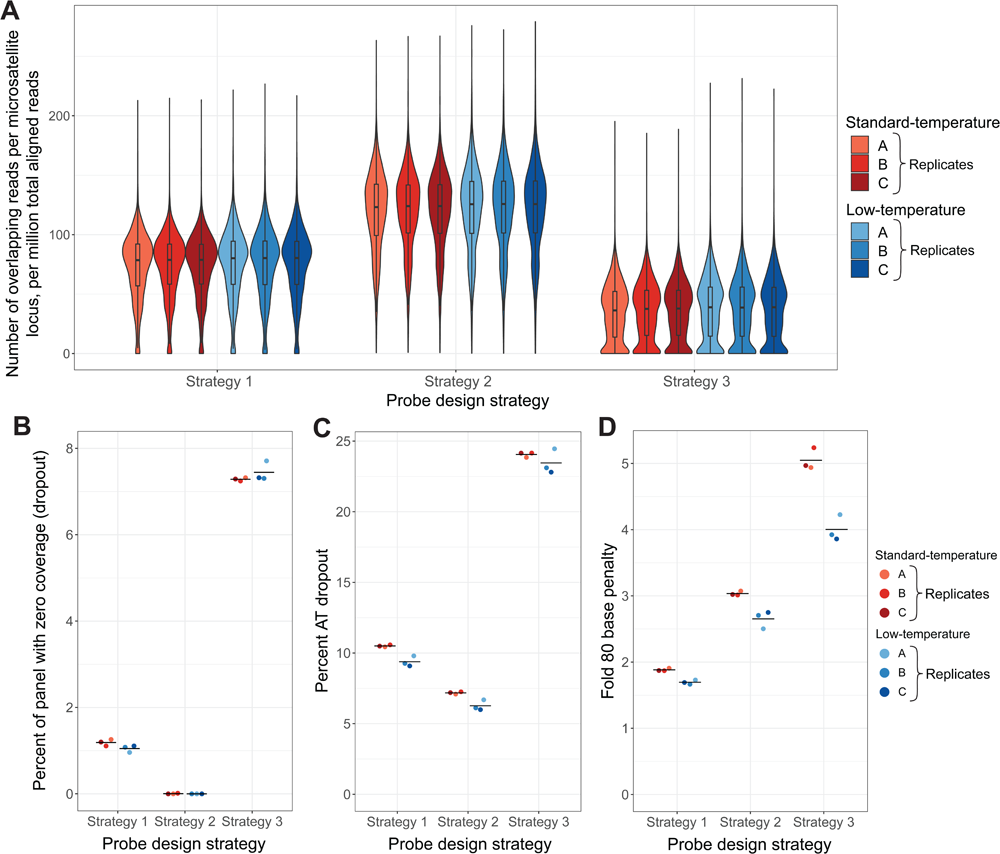
Hybridization capture quality metrics for different library amplification temperatures. **A.** Distribution of coverage across targeted microsatellite loci for each probe design strategy in both the standard (red) and low-temperature (blue) conditions. Normalized coverage is plotted as the [number of microsatellite-spanning reads at the locus] / [number of million aligned reads in the sample]. Box-and-whiskers show the first quartile, median, and third quartile of the distributions. Whiskers show 1.5 x the interquartile range. **B-D.** Hybridization capture quality metrics for low-temperature (blue, 3 technical replicates) and standard (red, 3 technical replicates) library amplification using the pilot capture panel, for each probe design strategy. **B.** Percent of panel with zero coverage. **C.** Percent AT dropout, which is calculated as the absolute value of the [percent of the target territory with ≤ 50% GC content] – [percent of aligned reads mapping to that territory]. High AT dropout indicates AT-rich sequences are being underrepresented in the final sequencing data relative to expectations due to capture biases (Zhou et al. 2021). Strategy 3 loci are likely more prone to dropout due to weak binding to probes, because only 60 bp of each probe binds to each flank instead of 120 bp probe binding in strategies 1 and 2. **D.** Fold 80 base penalty is a measure of capture uniformity and is calculated as the fold additional coverage needed to bring 80% of bases up to the mean coverage level. Greater uniformity is desired so that sequencing resources are not wasted by over-sequencing well-captured loci and by under-sequencing other loci. Strategy 2 has a higher fold 80 score than Strategy 1 because it has a wider distribution of coverage levels across loci, so under-covered loci require more additional coverage to reach the mean. Strategy 3’s even higher fold 80 base penalty is due to the larger number of loci with near-zero or zero coverage. Note, the standard-temperature samples shown in this figure are the same samples showin in Fig. 1C**-E**.

**Figure S6.**
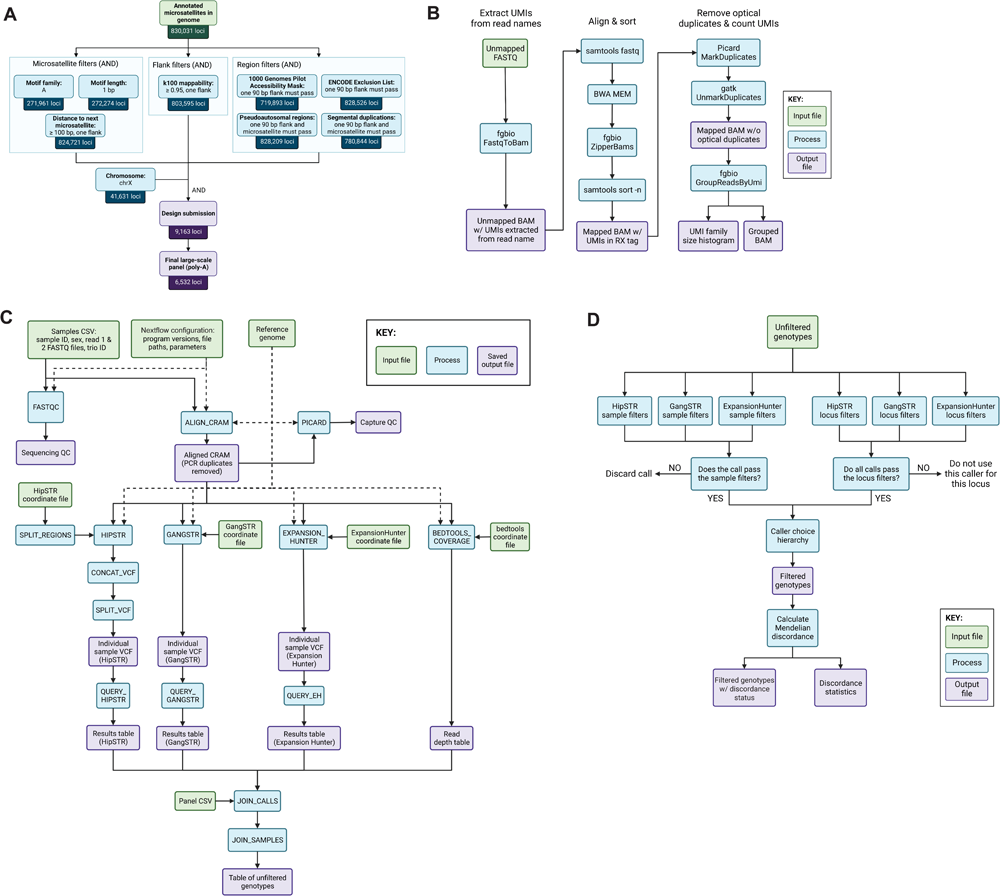
Selection of poly-A loci for the large-scale capture panel and schematic of the microsatellite genotyping pipeline, STREAM. **A.** Schematic of selection of poly-A loci for the large-scale microsatellite capture panel. All filters are applied with Boolean AND logic. **B.** Schematic of the computational pipeline used to calculate unique molecular identifier (UMI) duplication statistics. ‘w/’, with; ‘w/o’, without. **C,D.** Detailed schematics of the STREAM computational pipeline used to call genotypes of microsatellite loci with an ensemble approach (see **Methods** for full details). **C.** Stage 1 of STREAM checks sequencing quality, aligns reads to the genome, and generates microsatellite genotypes from HipSTR, GangSTR, and ExpansionHunter. **D.** Stage 2 of STREAM filters genotype calls based on various quality parameters, then selects for each locus a caller from which final genotypes are drawn based on quality parameters. Finally, the pipeline calculates the Mendelian discordance rate in family trios.

**Figure S7.**
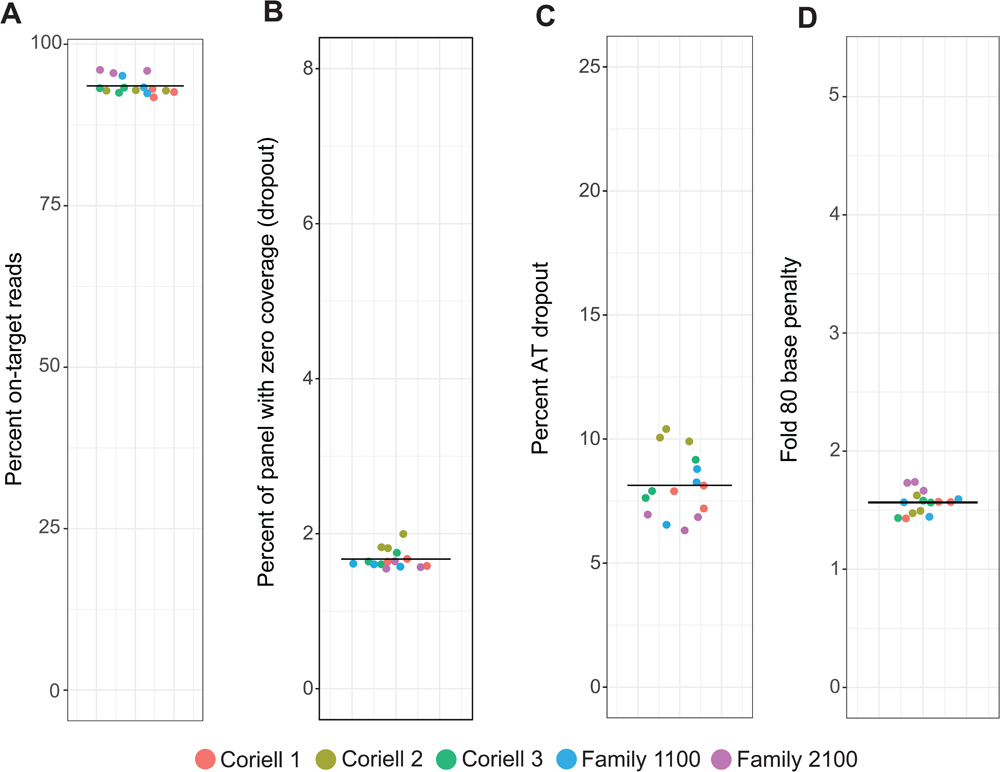
Hybridization capture quality metrics for the large-scale microsatellite panel. Large-scale microsatellite capture metrics for all family trio samples. Colors label family trios. **A.** Percent on-target reads. **B.** Percent of targets with zero coverage. **C.** Percent AT dropout. **D.** Fold 80 base penalty.

**Figure S8.**
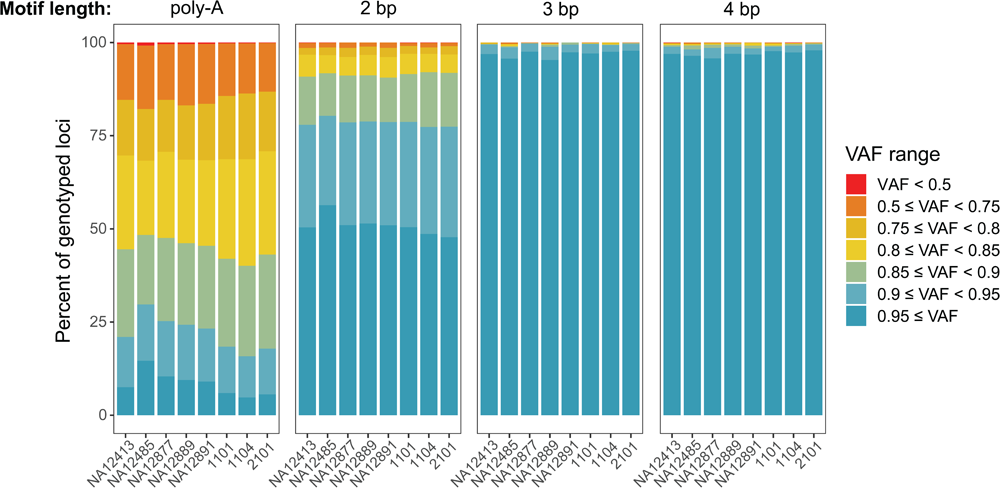
Microsatellite stutter analysis in the large-scale capture panel. Fraction of genotyped X-chromosome microsatellite loci in the large-scale panel in each variant allele fraction (VAF) range, with VAF calculated as in Fig. 2C. Only chromosome X loci from male individuals are included in the analysis to provide accurate stutter estimates, which is not feasible for bi-allelic loci. A VAF < 1 indicates stutter reads at the locus.

**Figure S9.**
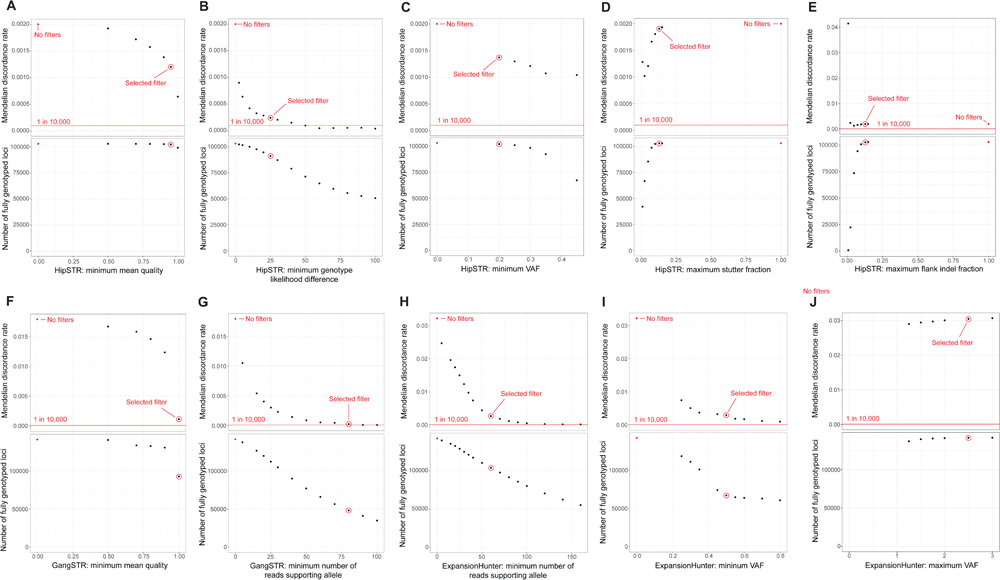
Optimization of microsatellite genotyping parameters. Effect of different quality filtering thresholds on the Mendelian discordance rate (top) and on the number of “fully genotyped” loci (i.e. loci with genotypes called in all family members) (bottom). Representative data shown here is for Family 1100 that was used for threshold optimization. During this optimization stage analyzing each individual parameter, only genotypes from the caller reporting that parameter were used and no secondary or tertiary caller was specified in the ensemble genotyping. Red dot: values without quality filtering. Red circle: threshold value chosen for final filtering settings. Red line: approximate expected microsatellite de novo mutation rate (i.e., Mendelian discordance rate) based on prior studies (Weber and Wong 1993; Huang et al. 2002; Sun et al. 2012; Kristmundsdottir et al. 2023). **A.** HipSTR: minimum mean quality score across all samples. **B.** HipSTR: minimum genotype likelihood difference (minimum likelihood difference between the reported genotype and the next most likely genotype). **C.** HipSTR: minimum variant allele fraction (VAF) of the called genotype. **D.** HipSTR: maximum fraction of reads with stutter. **E**. HipSTR: maximum fraction of reads with flank indels. **F.** GangSTR: minimum mean quality score across all samples. **G.** GangSTR: minimum number of reads supporting the allele. **H.** ExpansionHunter: minimum number of reads supporting the called allele. **I.** ExpansionHunter: minimum VAF of the called genotype. **J.** ExpansionHunter: maximum VAF of the called genotype (see **Methods** for an explanation of the need for this filter).

**Figure S10.**
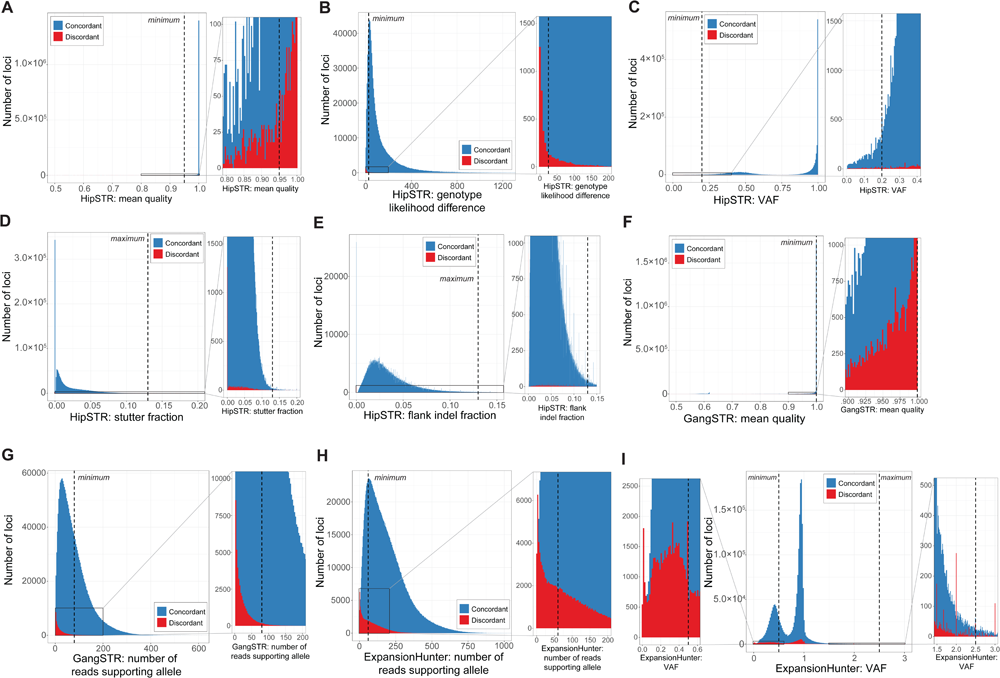
Distribution of filtering parameter values by concordance status. Overlayed histograms of parameter values across all loci for various filtering parameters. Histograms are generated from data from all 15 samples captured with the large-scale panel. Plots include only fully genotyped loci, as only these loci have concordance calls. Concordant calls are blue and discordant calls are red. The zoom plot shows the distribution of discordant calls near the final filtering threshold. Dashed line: chosen threshold in our final filter settings. **A.** HipSTR: mean quality score across all samples. **B.** HipSTR: genotype likelihood difference (minimum likelihood difference between the reported genotype and the next most likely genotype). **C.** HipSTR: variant allele frequency (VAF) of the called genotype. **D.** HipSTR: fraction of reads with stutter. **E**. HipSTR: fraction of reads with flank indels. **F.** GangSTR: mean quality score across all samples. **G.** GangSTR: number of reads supporting the allele. **H.** ExpansionHunter: number of reads supporting the allele. **I.** ExpansionHunter: VAF of the called genotype.

**Figure S11.**
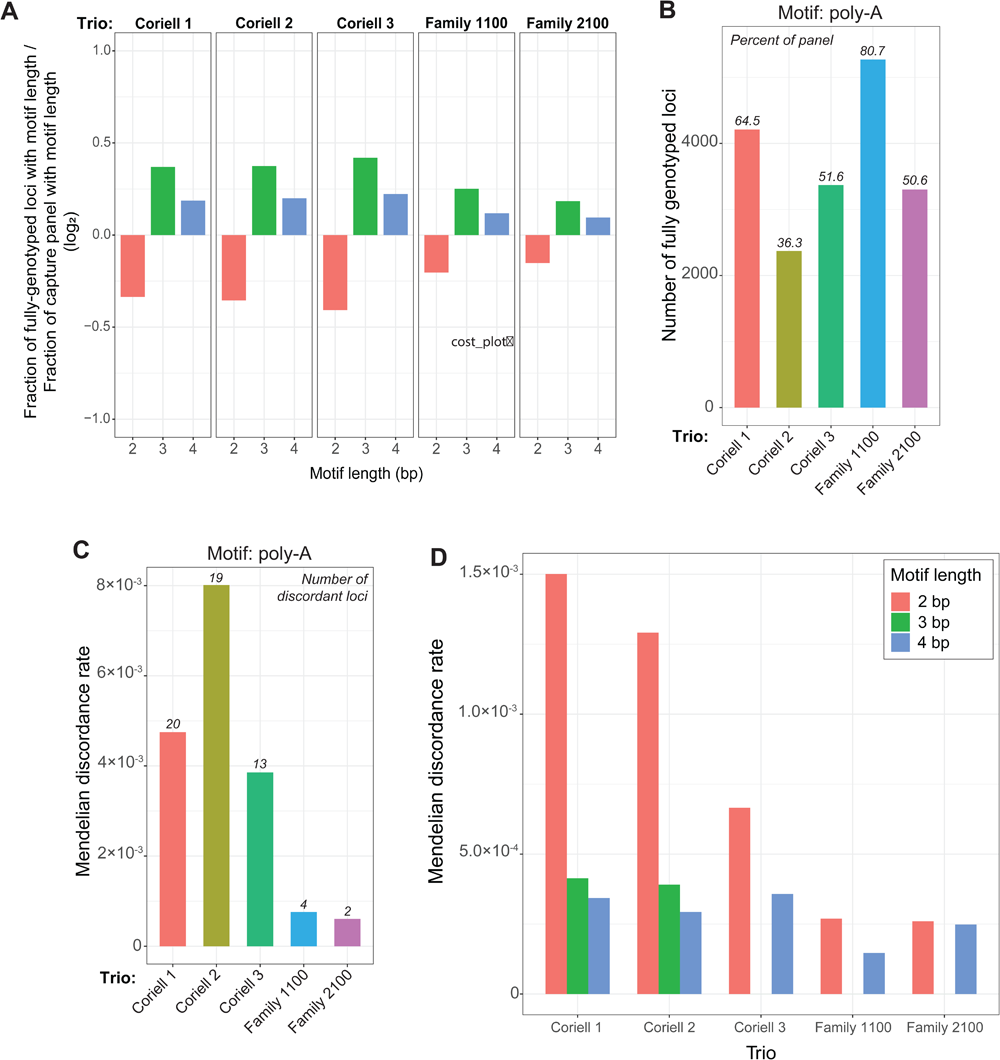
Genotyping and Mendelian discordance rates for different motif lengths. **A.** Log2 ratio of the fraction of fully genotyped loci that have 2, 3, and 4 bp motif lengths divided by the fraction of each respective motif length in the capture panel. **B.** Number of fully genotyped poly-A loci for each family trio; i.e. loci with genotypes called for all three members of the trio. Italics: fraction of capture panel that is fully genotyped. **C.** Mendelian discordance rate of fully genotyped poly-A loci in each family trio. Mendelian discordance rate = [number of discordant loci] / [number of fully genotyped loci]. Italics: number of discordant poly-A loci. **D.** Mendelian discordance rates for each motif length between 2 to 4 bp in each family trio.

**Figure S12.**
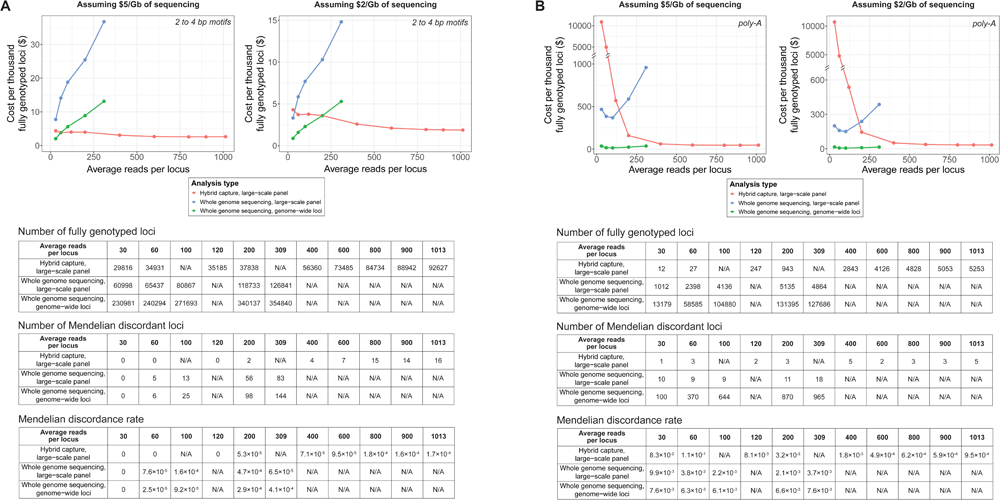
Cost-effective analysis of large-scale targeted capture versus whole-genome sequencing. **A,B.** Comparison of the cost per 1,000 fully genotyped loci (i.e., loci genotyped in all three members of the trio) for 2 to 4 bp motif loci (A) and poly-A loci (B) between hybridization capture with our large-scale panel, whole genome sequencing (WGS) with the loci from our large-scale panel, and WGS with a larger genome-wide list of 563,770 (395,725 loci with 2 to 4 bp motifs and 168,045 loci with a poly-A motif). The analysis is performed for different subsampled coverage levels (i.e., average reads per locus, which is calculated using all loci, including those that are not fully genotyped). See **Methods** for details. Tables list the number of fully genotyped loci, the number of Mendelian discordant loci, and the Mendelian discordance rates for each analysis method and coverage level. Mendelian discordance rates increase with read depth, because loci with non-discordant reference alleles are preferentially fully genotyped at lower read depths compared to loci with discordant non-reference loci.

Table S1. List of microsatellites targeted in the pilot capture panel

The list of microsatellites targeted in the pilot hybridization capture panel, which was used to profile technical replicates of NA12877 in both standard-and low-temperature amplification conditions. Each microsatellite is targeted by one of the three probe design strategies.

Table S2. Sample information and sequencing statistics for the pilot panel experiment

Sample information, sequencing statistics, and capture statistics for the pilot panel experiment.

Table S3. Comparison of stutter metrics using different library amplification temperatures

The percent of stutter reads (reads in the MALLREADS field of the HipSTR VCF that support a different number of repeat units than the called genotype) in loci of different motif lengths in standard-and low-temperature amplification conditions. We also calculate the percent of stutter reads that are shorter or longer than the called genotype and the percent of stutter reads where a single repeat unit was inserted or deleted. Statistics are the average across 3 technical replicates profiled with the pilot panel.

Table S4. List of microsatellites targeted in the large-scale capture panel

The list of microsatellites targeted in the large-scale hybridization capture panel, which was used on the family trio samples (**Table S5**).

Table S5. Family trio sample information and sequencing statistics

Sample information for the family trio samples profiled using the large-scale panel, with accompanying sequencing and capture quality statistics.

Table S6. Capillary electrophoresis validation primer sequences and results

The list of microsatellites for which capillary electrophoresis validation was performed, primer and amplicon information, and validation results from manual review in IGV (Integrative Genomics Viewer) and capillary electrophoresis.

